# YBEY is an essential biogenesis factor for mitochondrial ribosomes

**DOI:** 10.1101/2019.12.13.874362

**Authors:** Sabrina Summer, Anna Smirnova, Alessandro Gabriele, Ursula Toth, Fasemore Mandela, Konrad U. Förstner, Lauriane Kuhn, Johana Chicher, Philippe Hammann, Goran Mitulović, Nina Entelis, Ivan Tarassov, Walter Rossmanith, Alexandre Smirnov

## Abstract

Ribosome biogenesis requires numerous trans-acting factors, some of which are deeply conserved. In Bacteria, the endoribonuclease YbeY is believed to be involved in 16S rRNA 3’-end processing and its loss was associated with ribosomal abnormalities. In Eukarya, YBEY appears to generally localize to mitochondria (or chloroplasts). Here we show that the deletion of human YBEY results in a severe respiratory deficiency and morphologically abnormal mitochondria as an apparent consequence of impaired mitochondrial translation. Reduced stability of 12S rRNA and the deficiency of several proteins of the small ribosomal subunit in *YBEY* knockout cells pointed towards a defect in mitochondrial ribosome biogenesis. The specific interaction of mitoribosomal protein uS11m with YBEY suggests that the latter recruits uS11m to the nascent small subunit in its late assembly stage. This scenario shows similarities with final stages of cytosolic ribosome biogenesis, and may represent a late checkpoint before the mitoribosome engages in translation.

## INTRODUCTION

Ribosome biogenesis is a highly complex process that starts co-transcriptionally and includes ribosomal RNA processing, modification, and binding of ribosomal proteins (1). Each of these steps relies on specific factors, some of which are remarkably conserved. One such factor is the UPF0054 family protein YbeY found in all classified bacteria (2). Based on studies in various bacteria, YbeY has been implicated in ribosome maturation and quality control, with a particularly important role in small subunit (SSU) biogenesis (3-8), and post-transcriptional gene expression regulation (9-14). The deletion of *ybeY* is often lethal or associated with severe alterations of cellular metabolism and growth, indicating its indispensability for a wide variety of bacterial-type ribosomes (4,6,7,13-17).

Mechanistically, YbeY has been described as a metal-dependent endoribonuclease (5,12), and in some bacteria, *ybeY* mutants accumulate 16S rRNA with an unprocessed 3’ end (3,5,7,8). Therefore, YbeY was proposed to be the “missing” 3’ endoribonuclease required for 16S rRNA maturation to obtain the correct anti-Shine-Dalgarno sequence, which is needed for translation initiation on most bacterial mRNAs. However, this 16S rRNA 3’-misprocessing phenotype could equally be caused by the loss of a ribosome biogenesis factor that is not *per se* involved in rRNA cleavage (18), and so the precise role of YbeY in ribosome biogenesis remains unclear.

By carrying out an in-depth phylogenetic analysis, we found that YBEY is also conserved in many eukaryal lineages, including animals, plants, most stramenopiles and alveolates (Supplementary Figure 1). Indeed, YbeY of *Arabidopsis thaliana* was reported to be an essential ribosome biogenesis factor in chloroplasts, and its absence was associated with severe misprocessing of nearly all chloroplast rRNAs, resulting in deficiency of organellar translation, and hence, the absence of photosynthesis (16). Human YBEY, which shares 27% of identity with YbeY of the α-proteobacterium *Sinorhizobium meliloti* (15,19), has been predicted to localise in mitochondria (20), suggesting a role in human mitochondrial ribosome biogenesis. However, mitochondrial rRNAs are co-transcribed in a polycistronic precursor transcript with flanking tRNAs, and the mitochondrial tRNA processing enzymes RNase P and RNase Z are sufficient for their release (21-23). Moreover, mitochondrial mRNAs are leaderless and, therefore, do not rely on Shine-Dalgarno sequences for translation initiation (24). These considerations make an enzyme like YBEY apparently superfluous in the mitochondrial genetic system and raise the questions of why it has been retained in evolution and why, based on results of a recent genome-wide “death screen”, it seems to be required for life (25).

Here, we report a detailed characterisation of human YBEY and show that it is, indeed, an essential mitochondrial protein, required for mitochondrial translation and, therefore, cellular respiration. We show that it specifically interacts with the conserved mitochondrial chaperone p32 and mitoribosomal components and is crucial for the assembly of initiation-competent mitochondrial small subunits, apparently by recruiting the key ribosomal protein uS11m. This essential pathway, which may be conserved in other bacterial and bacteria-derived (i.e., mitochondria and plastids) genetic systems, shows striking parallels with the final steps of cytosolic small subunit maturation mediated by the adenylate kinase Fap7/hCINAP, suggesting that human cells use conceptually similar mechanisms to complete SSU assembly in the two translationally active compartments.

## MATERIAL AND METHODS

### Bacterial strains

*E. coli* strains used in this study (Table S1) are either BL21 Star (DE3) or Rosetta strains, adapted for recombinant protein production. For regular culturing, bacteria were grown at constant shaking at 200 rpm at 37°C in the standard liquid LB medium in the presence of appropriate antibiotics (in function of the hosted plasmids – see Table S1; Rosetta strains were routinely cultured in the presence of 34 µg/ml chloramphenicol; where needed, ampicillin and/or kanamycin were added at 100 µg/ml and 25 µg/ml, respectively).

### Human cell lines

293T-REx (Thermo Fischer Scientific), Flp-In T-REx 293 (Thermo Fischer Scientific), SAL001, HepG2 and HeLa cells (see Table S2 for the complete list of used cell lines) were cultured at 37°C, 7% CO_2_ in standard Dulbecco’s modified Eagle’s medium (DMEM) containing 4.5 g/l glucose supplemented with 10% fetal bovine serum. Medium was changed routinely every 48 h. For passaging, cells were washed with 1× phosphate buffered saline (PBS), resuspended in fresh medium and the required dilution was prepared. *YBEY* knockout cells were maintained in standard medium containing 1 mM pyruvate and 50 µg/ml uridine. Complemented *YBEY* knockout cells were grown in standard DMEM medium supplemented with 5 µg/ml blasticidin and 100 µg/ml zeocin.

For preparation of cell line stocks, confluent cells were harvested at 150*g* for 5 min. and resuspended in freezing medium (DMEM supplemented with 20% FBS and 10% dimethyl sulfoxide); 500 µl aliquots were frozen slowly at -80°C and stored in liquid nitrogen. Each batch of frozen cells was routinely tested for mycoplasma contamination.

All 293T-REx-derived cell lines in this study were authenticated by PCR, Sanger sequencing (for the *YBEY* gene), and western blotting (for the YBEY protein), as described in sections “CRISPR knockout cell line generation” and “*YBEY* knockout complementation”.

SAL001 cells and their corresponding parental control cell line (Flp-In T-REx 293) were induced with 0.1-0.2 mg/ml tetracycline to overexpress YBEY-3×FLAG 24 h before the experiment.

### CRISPR knockout cell line generation

The plasmid pX330 (26) was modified to generate pX330g containing a CMV promotor-driven EGFP expression cassette cloned in the NotI/Esp3I sites. pU6-gRNA1 was generated from pX330g by cleavage at the XbaI sites and re-ligation to delete the Cas9 expression cassette. For construction of pU6-YBEY-gRNA1 and pU6-YBEY-gRNA2 (from pU6-gRNA1) and pX330g-YBEY-gRNA3 and pX330g-YBEY-gRNA4 (from pX330g), the corresponding vector was digested with BbsI, dephosphorylated and used for ligations as previously described (26). Inserts were prepared by phosphorylation and annealing of the respective oligonucleotide pairs YBEY_gRNA_1f/YBEY_gRNA_1r, YBEY_gRNA_2f/YBEY_gRNA_2r, YBEY_gRNA_3f/YBEY_gRNA_3r and YBEY_gRNA_4f/YBEY_gRNA_4r (see Table S3 for the complete list of oligonucleotides used in this study). pPGK-puro was generated from pX260 (26) by cleavage at the EcoRI and NdeI sites to remove the Cas9 cassette, followed by blunt-ending with Klenow polymerase and re-ligation. All plasmids were verified by Sanger sequencing (primer pX260_1r).

For the generation of *YBEY*^+^ and *YBEY* knockout cell lines, subconfluent 293T-REx cells (2 cm^2^) were transfected with 0.2 µg of pU6-YBEY-gRNA1 and 0.4 of µg pX330g-YBEY-gRNA4 plasmids. To pre-select the successfully-transfected clones, the cells were co-transfected with 0.06 µg of pPGK-puro plasmid. The transfection mix, containing 1.5 µl of TurboFect (Thermo Fisher Scientific) and 0.66 µg of DNA in DMEM, was incubated for 15 min at room temperature and added to the cells. Twenty-four hours after transfection, the cells were transfered for two days in standard medium supplemented with 1 µg/ml puromycin. After pre-selection, the knockout cells were seeded at 1, 2, 4, or 8 cells per well in a 96-well plate in standard medium supplemented with home-made 30% conditioned medium. Cell clones were isolated and characterized by PCR with YBEY_6f/YBEY_6f primers, yielding a 486 bp product for the WT *YBEY* allele and 201 bp (Δ1), 31 bp (Δ2) and 130 bp (Δ3) for the disrupted alleles. As in the first round no homozygous *YBEY*-deletion clones were obtained, three *YBEY*^+^ clones were expanded and re-transfected, following the same protocol mentioned above, to generate knockout cells in a second round. *YBEY*^+^ clone 1 was transfected with 0.2 µg of pU6-YBEY-gRNA2 and 0.4 µg of pX330g-YBEY-gRNA3; *YBEY*^+^ clones 2 and 3 were re-transfected with 0.2 µg of pU6-YBEY-gRNA2 and 0.4 of µg pX330g-YBEY-gRNA4. Three knockout clones were finally isolated following the same procedure as described above and confirmed by PCR and western blotting.

### CRISPR knockout complementation

To construct the pcDNA4-YBEY plasmid, used for *YBEY* KO complementation with a WT *YBEY* allele, a PCR product generated with primers YBEY_3f and YBEY_7r, spanning the complete coding sequence of *YBEY* (167 aa, NM_058181), was inserted into the BamHI and XhoI sites of pcDNA4/TO. Plasmids with mutant *YBEY* alleles (pcDNA4-YBEY-R55A and pcDNA4-YBEY-H128A) were generated by QuickChange site-directed mutagenesis (Agilent) with the following pairs of primers, respectively: YBEY_R55A_1f/YBEY_R55A_1r and SAO00050/SAO0051. All plasmids were verified by Sanger sequencing (primers CMV_for and BGH_rev).

To generate *YBEY* knockout cells stably expressing WT YBEY or the YBEY mutant variants YBEY^R55A^ and YBEY^H128A^, subconfluent *YBEY* knockout cells of clone 2 and 3 (2 cm^2^) were transfected with ScaI-linearized pcDNA4-YBEY, pcDNA4-YBEY-R55A, or pcDNA4-YBEY-H128A, respectively. After 48 h, successfully-transfected cells were selected for 4 weeks using 5 µg/ml blasticidin and 100 µg/ml zeocin. Selected clones were tested for expression of YBEY by western blotting, and the identity of mutations was confirmed by PCR (the insert was amplified with primers CMV-for2 and BGH rev2 and sequenced with YBEY_3f and BGH-rev).

### Generation of a stable YBEY-3×FLAG cell line

The pSAP0006 plasmid to overexpress YBEY-3×FLAG was created by inserting the *YBEY* cDNA (amplified with primers SAO00029/SAO00030) into the ApaI/BspTI sites of pcDNA5 FRT/TO. Flp-In T-REx 293 cells were reverse-transfected with a mixture of 0.4 µg of pSAP006 and 3.6 µg of pOG44 with 10 µl of Lipofectamine 2000 (Thermo Fisher Scientific) in 2 ml of OptiMEM (Gibco) during 6 h. Medium was changed to EMEM, and the cells were left to propagate for 48 h. Then the cells were trypsinised, diluted and reseeded in the presence of 320 µg/ml hygromycin B Gold (InvivoGen). In 48 h, concentration of hygromycin was reduced to 160 µg/ml, and 18 days after transfection, individual clones were isolated and propagated in the presence of 160 µg/ml hygromycin. The clone used in this study is referred to as SAL001.

### siRNA-mediated gene knockdown

For uS11m silencing, 293T-REx cells were subjected to four sequential reverse transfections with 50 nM siRNA. For this, siRNA was complexed with Lipofectamine RNAiMAX (Thermo Fisher Scientific) according to the manufacturer’s protocol (e.g., for a 24-well plate, 1 µl Lipofectamine per 500 µl of OptiMEM was used) and incubated with cells for 6 h. Between transfections, cells were cultivated in DMEM during 48-72 h. Typically, cells transfected with uS11m-directed siRNAs show fast medium acidification after the fourth transfection, indicating a mitochondrial dysfunction.

### Transient complementation of *YBEY* KO

The pSAP0118 plasmid, representing a pcDNA5 FRT/TO variant with a weakened (Δ5) CMV promoter (27), was generated from pcDNA5 FRT/TO by QuikChange with SAO00217/SAO00218 primers. The same primers were used to derive the pSAP0109 plasmid from pSAP0006, permitting an attenuated YBEY-3×FLAG expression upon transfection. To construct an analogous uS11m-overexpressing plasmid (pSAP0119), the *MRPS11* cDNA was amplified with SAO00210/SAO00211, inserted in the BspTI/ApaI sites of pSAP0118, and the stop-codon was corrected by QuikChange with SAO00219/SAO00220. All plasmids were verified by Sanger sequencing (CMV_for and BGH_rev primers).

For transient transfection, *YBEY* KO cells (clones 2 and 3) were seeded in a 12-well plate. For complex formation, 1.6 µg of plasmid DNA was mixed with 4 µl of Lipofectamine 2000 in 1 ml of OptiMEM and added to the cells for 6 h. Then the medium was changed to DMEM. In 48 h, the transfection was repeated, and 72 h later, the cells were harvested with ice-cold PBS, lysed in 1× Laemmli buffer and analysed by 12% SDS-PAGE followed by western blotting.

### Western blotting

Protein samples in Laemmli buffer were resolved by SDS-PAGE, the proteins were transferred on an Amersham Protran western blotting nitrocellulose (GE Healthcare) or an Amersham Hybond P PVDF (GE Healthcare) membrane by semi-dry transfer. The membrane was blocked with 1× TBS containing 0.1% Tween-20 and 10% skimmed milk, and incubated with primary antibodies (see the Table S4 for the complete list of antibodies) diluted in 1× TBS, 0.1% Tween-20 for 1 h at room temperature or overnight at 4°C. The membrane was washed with 1× TBS, 0.1% Tween-20 and incubated with a dilution of the corresponding HRP-coupled secondary antibody for 30 min, followed by another round of washing. The chemiluminescent signal was visualised with the SuperSignal West Pico PLUS (Thermo Fisher Scientific), or Westar Sun (Cyanagen), or ECL Select Western blotting (GE Healthcare) chemiluminescent substrate on ChemiDoc Touch (Bio-Rad) or G-Box (Syngene) and analysed with Image Lab (v. 5.2.1).

### Northern blotting

RNA samples mixed 1:1 with the denaturing gel loading buffer (0.025% SDS, 18 mM EDTA, 0.025% bromophenol blue, 0.025% xylene cyanol in deionised formamide) and boiled for 5 min at 95°C were separated on 6-8% polyacrylamide/7 M Urea denaturing gel in 1× TBE and transferred onto an Amersham HybondN+ membrane (GE Healthcare). RNA was UV-crosslinked to the membrane. Pre-hybridization was performed for 30 min at 65°C in 6× SSC containing 5× Denhardt’s solution and 0.2% SDS. The membrane was incubated overnight with a 5’-^32^P-labelled probe in 3× SSC, 0.1% SDS, 0.5× TE, 0.5 M NaCl, 5× Denhardt’s solution at 42°C with continuous rotation. After hybridization, the membrane was washed with 5× SSC, 0.1% SDS, dried, and exposed with a Phosphorimager plate. The radioactive signal was visualised on Typhoon Trio (GE Healthcare) and analysed with ImageQuant TL (v. 7.0, GE Healthcare). For re-probing, membranes were stripped in stripping buffer (1% SDS, 0.1x SSC, 40 mM Tris-HCl, pH 7.6) at 80°C three times for 10 minutes and washed once in 2× SSC.

### Subcellular and mitochondrial fractionation

For subcellular fractionation, confluent 293T-REx cells (25 cm^2^) were harvested and nuclei were isolated from 5×10^6^ cells with the Qproteome Cell Compartment Kit (QIAGEN), according to the manufacturer’s instructions. Lysates were incubated for 30 min on an end-over shaker at 4°C and centrifuged at 6,000*g* at 4°C.

Mitochondria were prepared from 1×10^7^ 293T-REx cells. Cells were washed twice with 1× PBS and resuspended in 400 µl of chilled RSB buffer (10 mM Tris-HCl, pH 7.6, 10 mM NaCl, 1.5 mM CaCl_2_). Cells were incubated for 15 min on ice, homogenized with a 27G needle on a 1 ml syringe, and 1 volume of chilled MS buffer (420 mM mannitol, 140 mM sucrose, 10 mM Tris-HCl, pH 7.6, 5 mM EDTA) was added. Cell debris was pelleted by centrifugation at 4°C as follows: 2 min at 500*g*, twice 2 min at 1000*g*, with transferring the supernatants to fresh tubes after each centrifugation. Mitochondria were pelleted at 10,000*g* at 4 °C for 8 min, washed once with chilled M3 buffer (210 mM mannitol, 70 mM sucrose, 20 mM Tris-HCl, pH 7.6, 10 mM KCl, 6 mM EDTA) supplemented with 1 mM DTT, 0.1% proteinase inhibitor cocktail (Roche) and incubated for 10 min on ice in 100 µl of chilled M3 buffer supplemented with 1 mM DTT, 0.1% proteinase inhibitor cocktail and 0.02% digitonin. Mitochondrial pellets were washed once with chilled M3 buffer supplemented with 1 mM DTT, 0.1% proteinase inhibitor cocktail. Total cell, mitochondrial and nuclei pellets of 293T-REx cells were resuspended in 1× PBS and an equivalent of 5×10^5^ cells was resolved by 15% SDS-PAGE. Purity of the fractions and the presence of YBEY in the cellular compartments were analyzed by western blotting.

Submitochondrial fractionation was performed as described in (28). Briefly, 225 cm^2^ of SAL001 cells were grown to 80-90% confluency and induced with 0.2 mg/ml tetracycline for 24 h. All subsequent manipulations were carried out at 4°C. The cells were resuspended in 1.5 ml of Breakage buffer (0.6 M sorbitol, 10 mM HEPES-KOH, pH 7.5, 1 mM EDTA) and disrupted with a syringe (26G×25 mm, 20 strokes). After two low-speed centrifugations (600*g* and 1,000*g*) for 10 min to remove cell debris and nuclei, the mitochondria-rich fraction was collected by high-speed centrifugation at 14,000*g* for 20 min. The pellet was resuspended in 0.5 ml of Breakage buffer, split in three equal aliquots and centrifuged for another 10 min at 14,000*g*. The supernatants were discarded. The pellets were resuspended at 0.5 mg/ml (mitochondrial protein measured by Bradford assay, Roti-Nanoquant, Carl Roth) in one of the following buffers: (i) Breakage buffer; (ii) “Swelling” buffer (10 mM HEPES-KOH, pH 7.5, 1 mM EDTA); (iii) Lysis buffer (10 mM HEPES-KOH, pH 7.5, 1 mM EDTA, 0.5% n-dodecyl-β-D-maltoside). The latter sample was additionally lysed with a syringe (10 strokes) to ensure complete membrane solubilisation. Each aliquot was then split in two equal portions, one of which was treated with 50 µg/ml proteinase K. The samples were incubated on ice for 20 min, added 1 mM PMSF and 1/4 volume of 100% trichloroacetic acid (prepared by dissolving 5 g of TCA in 3.5 ml of water), then incubated on ice for another 10 min. Upon a 10 min centrifugation at 14,000*g*, the precipitates were washed twice with ice-cold acetone and dried overnight on bench. They were then resuspended in 100 µl of 1×Laemmli buffer, heated for 5 min at 80°C and sonicated. For analysis, 10 µl of each sample was resolved by 15% SDS-PAGE followed by western blotting.

### Microscopy and image analysis

The pSAP0123 plasmid to overexpress YBEY-FLAG was generated from pSAP0006 by QuikChange with primers SAO0225/SAO0226.

For confocal microscopy, cells were seeded on an 8-well Nunc Lab-Tek slides (Thermo Fisher Scientific) 24 to 48 h prior the experiment. For Flp-In T-REx 293-derived cell lines, the slide was covered with 0.1 % gelatine to improve cell adhesion. For this the slide was incubated with gelatine for 30 min at 37°C at 5% CO_2_, then the excess of gelatine was washed out with MilliQ and the slide was dried. For transient transfection of cells with pSAP0123, 150 ng of plasmid DNA was mixed with 0.5 µl of Lipofectamine 2000 in 300 µl of OptiMEM and incubated with adherent cells during 4-6 h. After that, medium was changed to EMEM. In 24 h post-transfection, cells were fixed by incubation during 12 min at 37°C with 3% formaldehyde solution diluted with the DMEM medium (4% paraformaldehyde dissolved in PBS by heating at 60°C and adjusted to 3% with DMEM). For immunostaining, the cells were permeabilised with 0.3 % Triton X-100 in 1× PBS for 10 min at room temperature. After blocking with 5% bovine serum albumin (BSA) in 1× PBS for 30 min at room temperature, samples were incubated for 1 to 3 h with primary antibodies diluted in the blocking buffer, then with secondary antibodies conjugated with respective fluorophores. For nuclear counterstaining, 1 µg/ml DAPI in 1× PBS was applied for 5 min at room temperature. Each step alternated with five washes with 1× PBS. Samples were imaged on LSM700 or LSM780 microscopes (Carl Zeiss) under 63×/1.4 oil objective in Vectashield (Vector Laboratories) or Prolong Gold (Invitrogen) mounting media. Fluorescence intensity profiles were measured in ImageJ (29). For estimation of the uS11m amount in different cell lines, images were segmented by global Li thresholding method (30), then mean fluorescence intensity was calculated in ImageJ.

Simultaneous RNA smFISH and protein immunostaining were done according to the manufacturer’s protocol for the ViewRNA ISH Cell assay kit (Thermo Fisher Scientific). For RNA smFISH, the branched DNA technology was used (31): a gene-specific oligonucleotide target probe set containing 20 to 40 probe pairs binds to the target RNA sequence (see the complete list of smFISH probes in Table S3), and signal amplification is achieved through a series of sequential hybridization steps with pre-amplifier, amplifier and finally fluorophore-labelled oligonucleotides. For the analysis of RNA abundance in different cell lines, images of the labelled RNA and the mitochondrial reference protein were segmented and subcellular shapes were quantified with the Squassh method (32) in the MosaicSuit in Fiji (33). The obtained mean size of objects representing individual RNAs or groups of RNAs was multiplied by the number of the objects in each frame for estimation of total abundance of RNA. This parameter was then normalized to the total abundance of the mitochondrial reference protein (TOMM20 or mL38) obtained by the same procedure.

For FLIM-FRET analyses, SAL001 cells were immunolabelled, as described above, with primary and secondary antibodies coupled with synthetic fluorophores that form a FRET pair (34,35). The less abundant protein was chosen as fluorescence donor and labelled with Alexa Fluor 488; the more abundant acceptor protein was labelled with Alexa Fluor 555. For testing the YBEY-p32 interaction, YBEY was labelled with YBEY antibodies, in all other cases with FLAG antibodies. Camera-based lifetime detection of donor fluorescence was performed on a widefield microscope FLIM Nikon TE2000 operated in the frequency domain. Before each experiment, the system was calibrated at the pixel level with the reference fluorophore fluorescein with known fluorescence lifetime of 4 ns. The number of analysed frames is shown on Figure 5B.

**Figure 1:**
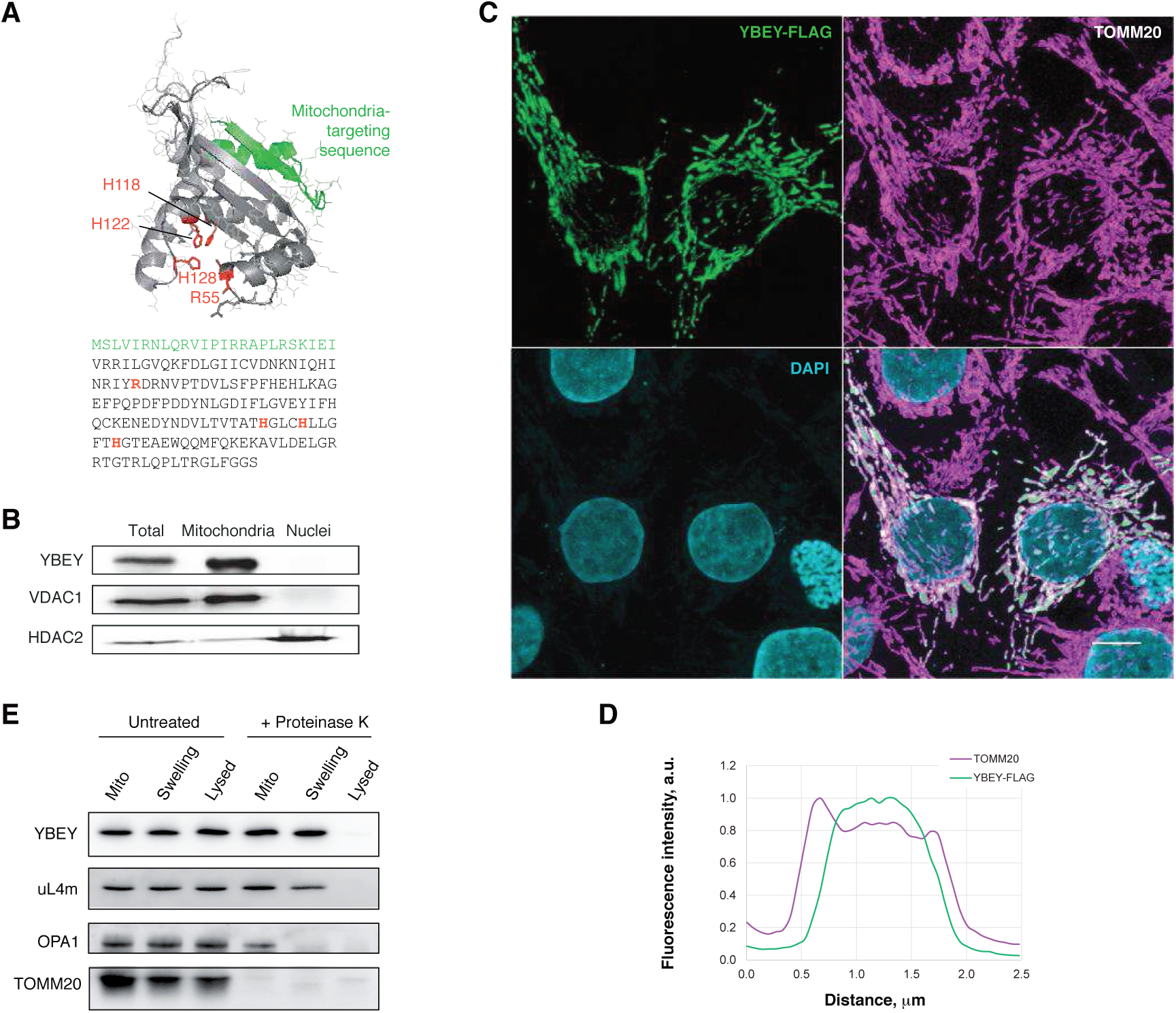
Human YBEY is a *bona fide* mitochondrial protein. **(A)** The structural model (RaptorX) and the sequence of human YBEY. The predicted mitochondria-targeting peptide (TargetP) is highlighted in green. The diagnostic histidine triad of the UPF0054 family and the highly conserved R55 residue, mutated in this study, are shown in red. **(B)** Subcellular fractionation of HEK293T-REx cells supports a mitochondrial localisation of endogenous YBEY. VDAC1 and HDAC2 are a mitochondrial and a nuclear marker, respectively. **(C)** Immunostaining of transiently transfected HepG2 cells expressing YBEY-FLAG. TOMM20 is used as mitochondrial marker. Scale bar, 10 µm. **(D)** Representative fluorescence intensity profiles of TOMM20 and YBEY-FLAG across a mitochondrion, derived from the experiment shown in (C), indicate the accumulation of YBEY in the interior space of mitochondria. **(E)** Submitochondrial localisation of the YBEY-3×FLAG protein. Crude mitochondria (“Mito”), or mitochondria with the outer membrane ruptured by hypotonic swelling (“Swelling”), or mitochondria lysed with 0.5% β-dodecyl maltoside (“Lysed”) were treated (or not) with proteinase K and analysed by western blotting.

**Figure 2:**
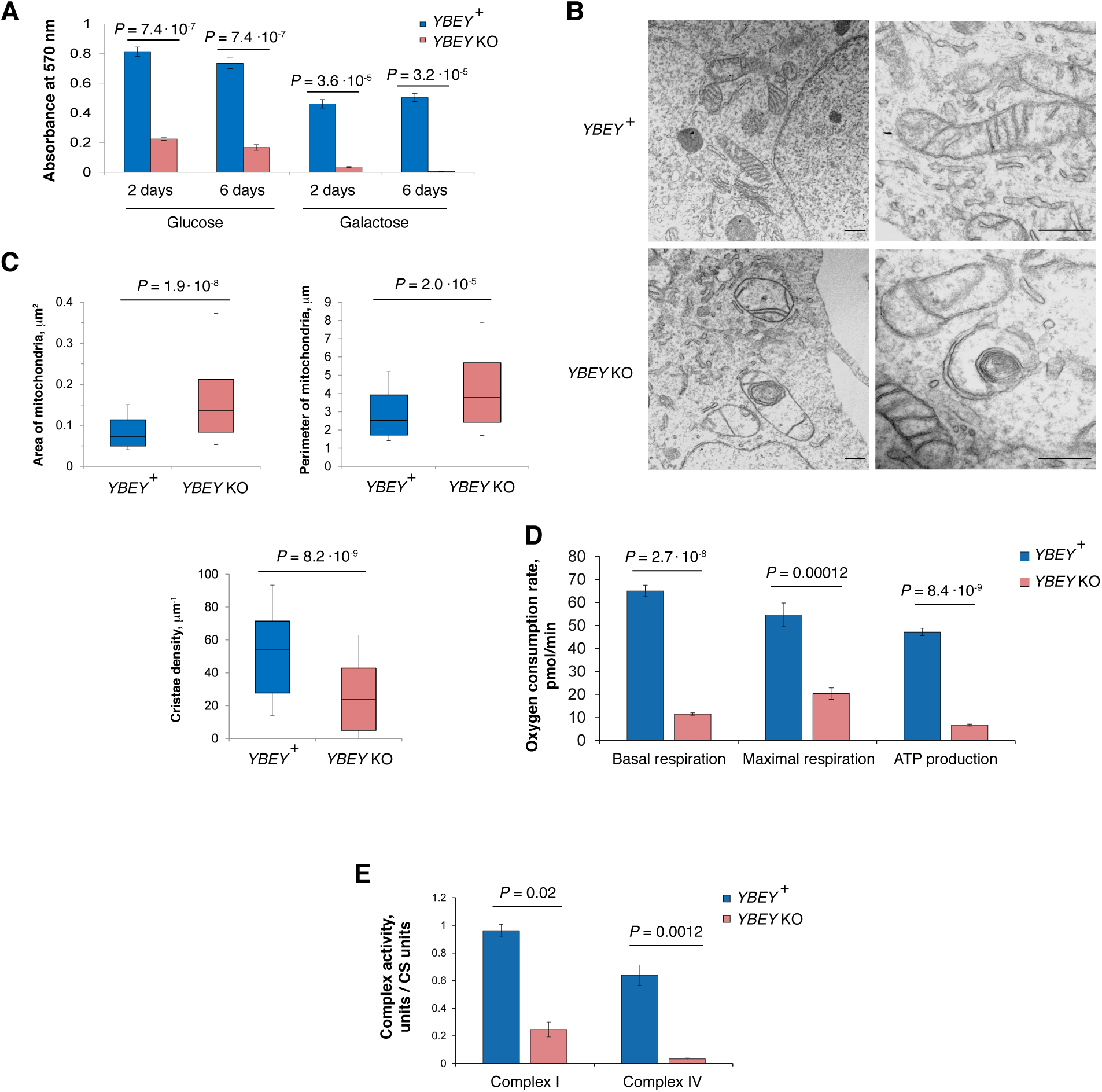
YBEY is required for normal mitochondrial morphology and respiration. **(A)** MTT assay for cell proliferation reveals a slower growth of *YBEY* KO cells in a glucose-containing medium and nearly no growth when galactose is used as carbon source. Means ± SEM for *n* = 12 are shown; *P*-values, two-tailed Mann-Whitney test. **(B)** Transmission electron microscopy of HEK293T-REx cells shows enlarged mitochondria with disorganised cristae systems in *YBEY* KO cells. A group of mitochondria (left) and individual organelles (right) are shown. Scale bars, 500 nm. **(C)** Quantitative analysis of mitochondrial morphology from images like in (B). Medians, interquartile ranges, 10^th^ and 90^th^ percentiles are shown. For *YBEY* ^+^ and *YBEY* KO cells, *n* = 116 and 88 mitochondria were analysed, respectively; *P*-values, two-tailed Mann-Whitney test. **(D)** Respiration phenotypes of *YBEY* KO cells, as assessed by oxygen consumption rate measurement with the Seahorse Mito Stress test. Means ± SEM for *n* = 8 (*YBEY* ^+^) and *n* = 7 (*YBEY* KO) are shown; *P*-values, two-tailed Welch’s test. **(E)** YBEY loss results in significantly decreased activities of respiratory complexes I and IV. Means ± SEM (*n* = 3) of citrate synthase (CS)-normalized activities are shown; *P*-values, two-tailed Welch’s test.

**Figure 3:**
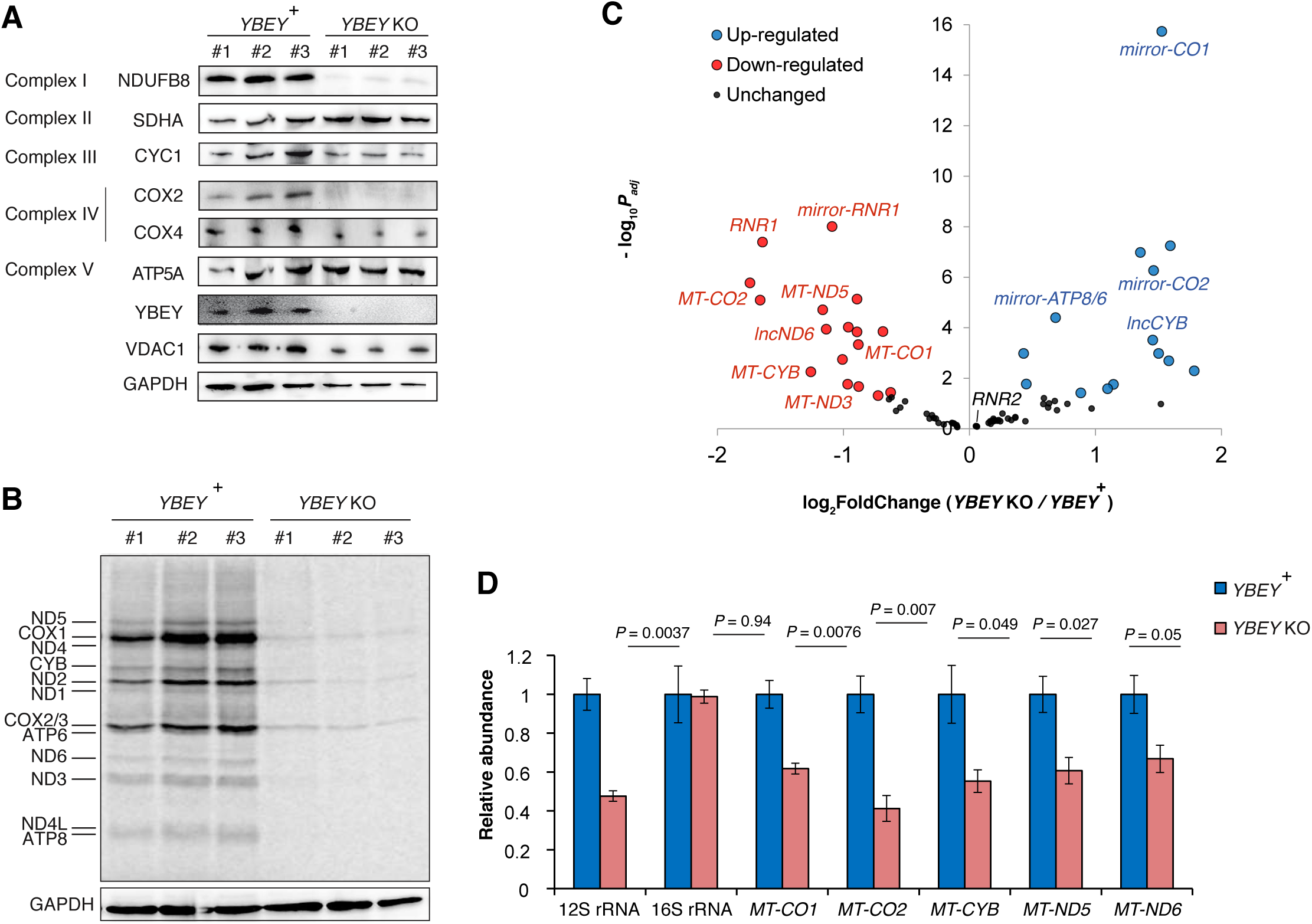
YBEY is essential for mitochondrial gene expression. **(A)** Western blot analysis of select respiratory chain subunits from *YBEY* KO cells shows selective depletion of the mtDNA-encoded COX2 protein and of the nucleus-encoded NDUFB8 subunit. VDAC1 and GAPDH are shown as loading controls. **(B)** Metabolic [^35^S]-methionine labelling of mitochondrial translation products reveals a nearly complete shutdown of protein synthesis in *YBEY* KO mitochondria. **(C)** Volcano plot summarising the gene expression changes between *YBEY* ^+^ and *YBEY* KO mitochondria as assessed by RNA-Seq. Select loci, further discussed in the text, are highlighted. See also Table S5. **(D)** RT-qPCR measurement of select mitochondrial RNAs in *YBEY*^+^ and *YBEY* KO cells. Means ± SEM for *n* = 3 are shown; *P*-values, two-tailed t-test.

**Figure 4:**
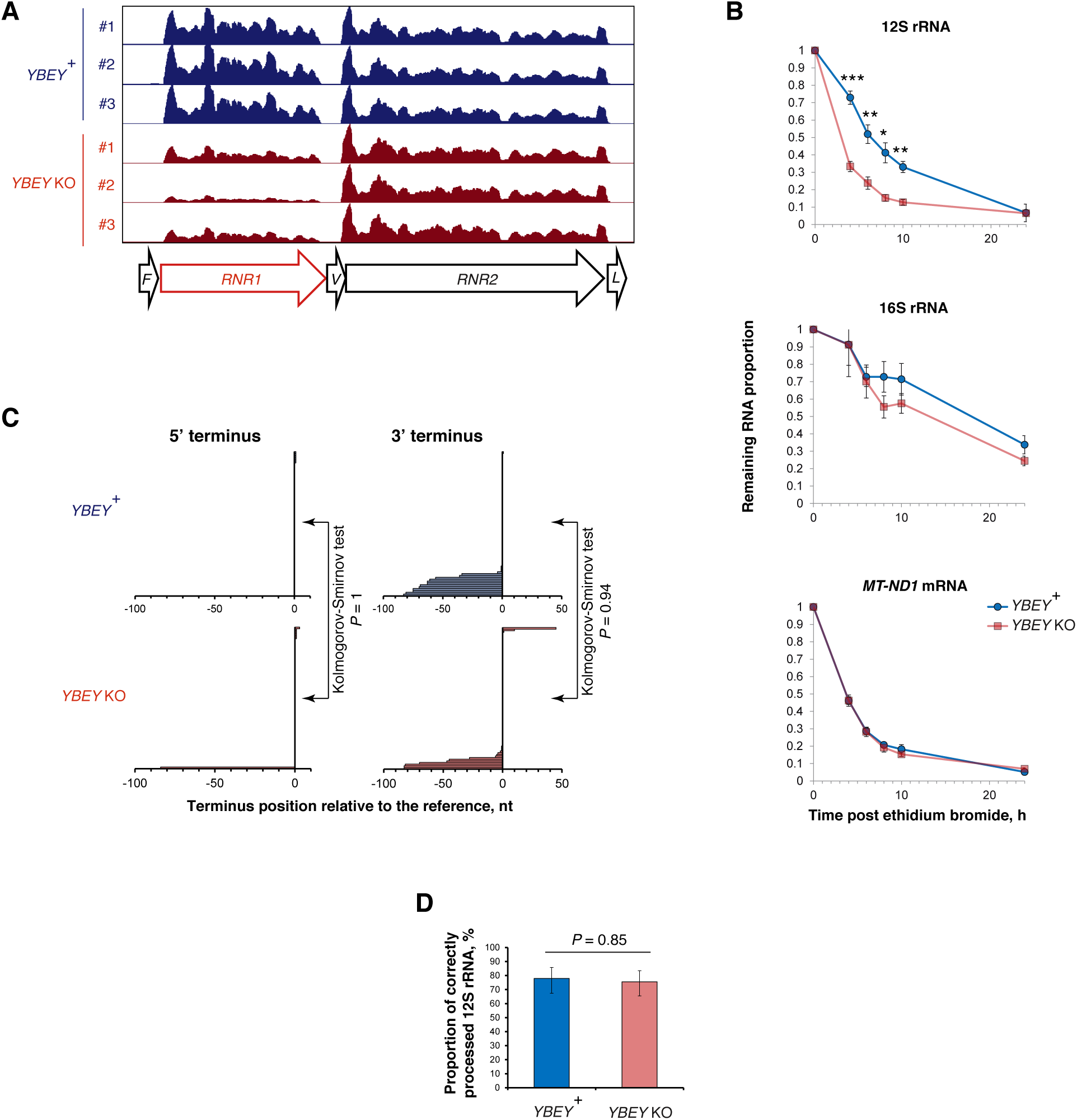
12S rRNA is destabilised, yet correctly processed in the absence of YBEY. **(A)** Snapshot of the mitochondrial rRNA locus showing a select downregulation of 12S rRNA in the absence of YBEY, as assessed by RNA-seq. tRNA genes are labelled with the single-letter code. **(B)** The half-life of 12S rRNA is significantly decreased in *YBEY* KO cells. RNA samples were collected at various time points after arresting mitochondrial transcription with ethidium bromide and analysed by RT-qPCR. Means ± SEM for *n* = 4 (*YBEY* ^+^) or *n* = 9 (*YBEY* KO) are shown; *P*-values, two-tailed Welch’s test: \**P* = 0.013, \*\**P* < 0.005, \*\*\**P* = 7.7 · 10^−5^. **(C)** cRT-PCR analysis of the 5’- and 3’-termini of 12S rRNA in *YBEY* ^+^ and *YBEY* KO cells. *n* = 77 plasmid clones from two independent *YBEY*^+^ cell lines and *n* = 86 clones from two independent *YBEY* KO cell lines are aligned with respect to the reference positions, the majority of clones showing the correct processing on both sides. **(D)** *YBEY*^+^ and *YBEY* KO cells have indistinguishable proportions of correctly processed 12S rRNA molecules. Proportions and 95% CIs are shown; based on the data in (C); *P*-value, Fisher’s exact test.

**Figure 5:**
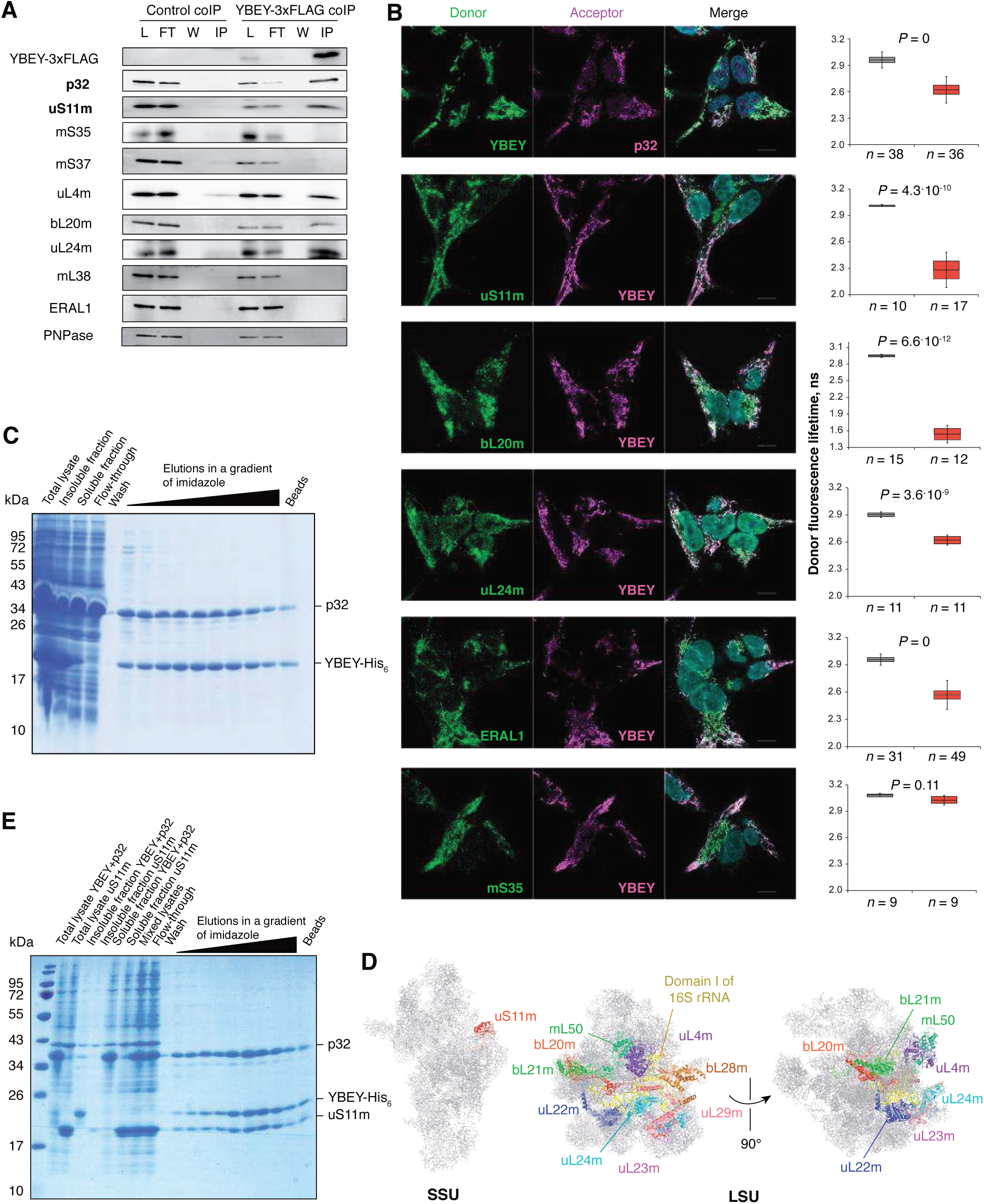
YBEY interacts with p32 and a distinct set of mitoribosomal proteins. **(A)** Immunoprecipitation of YBEY-3×FLAG from mitochondrial lysates copurifies p32 and select mitoribosomal proteins. Western blot analysis of lysate (L), flow-through (FT), wash (W) and immunoprecipitate (IP) fractions for WT control (not expressing FLAG-tagged baits) and stable YBEY-3×FLAG-expressing HEK293T-REx cells is shown. The particularly highly enriched p32 and uS11m proteins are set in bold. See also Table S6. **(B)** Indirect FLIM-FRET analysis of *in situ* interactions between YBEY and select mitochondrial proteins in HEK293T-REx cells. Representative images of staining of the corresponding partners with fluorescent antibodies forming a FRET couple are shown along with donor fluorescence lifetime measurements when the donor alone (grey boxes) or together with the acceptor (red boxes) is labelled. The boxes show the means and the 95% CI, the bars are SD. A significant decrease in the donor fluorescence lifetime in the presence of the acceptor (Bonferroni-corrected two-tailed Welch’s test) is indicative of FRET between the two and, therefore, of a tight spatial proximity of the interactors. On all microscopy images, the donor is labelled in green and the acceptor in magenta. mS35 is an example of mitoribosomal protein that does not interact with YBEY. **(C)** Coexpression of human YBEY-His_6_ with tagless p32 in *E. coli* results in partial solubilisation of the former and stoichiometric copurification of both on a Ni-agarose column. **(D)** Mapping of the mitoribosomal proteins copurifying with YBEY on the structure of the mammalian mitoribosome. Note the spatial clustering of the enriched LSU proteins around domain I of 16S rRNA. **(E)** A stable and stoichiometric ternary complex copurifies on Ni-agarose beads from mixed lysates of YBEY-His_6_/p32 and uS11m-expressing *E. coli* cells. See Supplementary Figure 9C for a control purification.

Transmission electron microscopy was performed at the Plateforme Imagerie In Vitro, CNRS, UPS3156, University of Strasbourg. Briefly, cells were washed with 1× PBS and fixed with 3% glutaraldehyde in 0.1 M cacodylate buffer. Then samples were dehydrated in a graded ethanol series, treated with propylene oxide, embedded in araldite resin, cut and mounted on copper grids. Ultrasections were contrasted with uranyl acetate. Specimens were examined with a Hitachi H 7500 microscope equipped with an AMT Hamamatsu digital camera. For quantitative analysis, mitochondria were outlined manually and their area and perimeter determined in ImageJ. The cristae numbers were counted manually and normalized to the mitochondrial area.

### MTT cell proliferation assay

Confluent cells were resuspended and seeded at 1×10^4^ cells (for the 2-days time point) and 6×10^2^ (for the 6-days time point) per well on a 96-well plate in glucose medium (glucose-free DMEM supplemented with 10% FBS, 2 mM glutamine and 5 mM glucose) or in galactose medium (glucose-free DMEM supplemented with 10% FBS, 2 mM glutamine and 5 mM galactose). For each time point and treatment, duplicates were prepared. After 2 days and 6 days of growth, respectively, 0.5 mg/ml thiazolyl blue tetrazolium bromide (MTT) was added to the medium. Cells were incubated for 2 h and 100 µl of isopropanol was added to each well. The absorbance was measured at 570 nm over 1 min with the Enspire multimode plate reader (Perkin Elmer). For all values the blank mean was subtracted.

### Seahorse respiration phenotype measurements

Cells were cultivated standardly in the DMEM medium, dissociated with a 0.25% trypsin-EDTA solution, and their amount was estimated by Sceptor Handled Automated Cell Counter (Millipore). Then the cells were seeded in the DMEM medium on Seahorse XF96 V3 PS cell culture microplates (Seahorse XFe96 FluxPak mini, Agilent) coated with 0.1 % gelatine 24 h before the experiment. To achieve measurable OCR, 10,000 cells (*YBEY*^+^ and complemented *YBEY* KO cell lines) or 30,000 cells (uncomplemented *YBEY* KO) per well were seeded. The Seahorse XF Cell Mito Stress test was performed on a Seahorse XFe96 Analyzer (Agilent) in the XF Base Medium (Agilent) supplemented with 25 mM glucose, 4 mM glutamine and 1 mM pyruvate (pH 7.4 adjusted at 37°C with 5 M NaOH). Compounds from the Seahorse XF Cell Mito Stress Test Kit (Agilent) were used in the following order and concentrations: 2 µM oligomycin, 2 µM FCCP, 0.5 µM rotenone/antimycin A. Basal respiration was measured as the difference between the OCR before the first injection and the non-mitochondrial respiration rate. ATP production was measured as the difference between the OCR before the oligomycin injection and the minimum OCR after the oligomycin injection. Maximal respiration is the difference between the maximum OCR after the FCCP injection and non-mitochondrial respiration. The latter is defined as the minimum OCR after the rotenone/antimycin A injection. ECAR was measured in the absence of additives. The data were analysed in Wave Desktop and Controller 2.6 (Agilent Technologies).

### Respiratory complex activity assays

Cells (75 cm^2^) were harvested and washed once with 1× PBS; 1×10^7^ cells were used for mitochondrial preparation by differential centrifugation, as described above. The initial mitochondrial pellets were frozen in liquid nitrogen and stored at -80°C until measurement. Mitochondria from 1×10^7^ cells were resuspended in 100 µl of SETH buffer (250 mM sucrose, 10 mM Tris-HCl, pH 7.4, 2 mM EDTA) and broken by two freeze-thaw cycles of freezing in liquid nitrogen and thawing on ice. Mitochondrial lysates were gently resuspended before the use in activity measurements.

For spectrophotometric measurements a UV-Vis spectrophotometer (Shimadzu UV-1800) was used. Complex I activity was determined by measuring the rotenone-sensitive NADH oxidation at 334 nm at 30 °C with the use of decylubiquinone as electron acceptor. For this, 30 µl of mitochondrial lysate was used in a total reaction volume of 750 µl. The reaction was performed in 25 mM potassium phosphate buffer, pH 7.4, containing 2.8 mg/ml BSA, 5 mM MgCl_2_, 0.2 mM NADH_2_, 2 mM KCN, 0.1 mM decylubiquinone and 4 µM antimycin A. Before adding the sample, the assay buffer was incubated for 5 min at 30°C and background absorbance was followed for 2 min. The sample was added and absorbance measured for 6 min. Rotenone (0.02 mM) was added to inhibit complex I activity, and rotenone-insensitive absorbance was measured for 4 min. The complex activity (units/l) was calculated as follows:

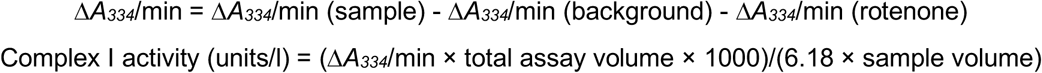

Complex IV activity (via oxidation of reduced cytochrome c (II)) was measured at 550 nm at 30°C in the assay buffer containing 85 mM HEPES-NaOH, pH 7.2, and 90 µM reduced cytochrome c (II) (Sigma-Aldrich). For reaction, 10 µl of mitochondrial lysate were used in a total volume of 750 µl. After 5 min pre-incubation, the sample was added and absorbance (background sample) at 550 nm was followed for 8 min at 30°C. Potassium hexacyanoferrat (III) was added in a final concentration 64.6 µM and the absorbance was followed for another 3 min at 30°C. The Complex IV activity (units/l) was calculated as follows:

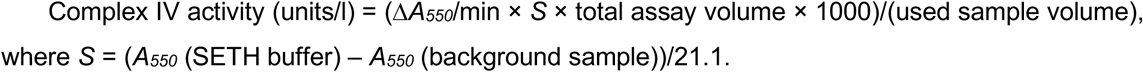

Activity of citrate synthase (CS) was determined by measurement of reduction of acetyl-coenzyme A (Ac-CoA) at 412 nm at 37°C. For this, 5 µl of mitochondrial lysate was added to 750 µl of the reaction buffer containing 0.1% Triton X-100, 0.1 mM 5,5′-Dithiobis(2-nitrobenzoic acid) and 0.1235 mM Ac-CoA. The sample was added to the buffer and the background absorbance was measured for 2 min. Then 0.5 mM of freshly prepared oxaloacetic acid was added and the absorbance was measured for another 2 min. Citrate synthase activity (units/l) was calculated as follows:

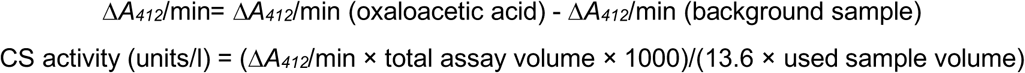

OXPHOS complex I and IV activities were normalized to CS activity.

### Mitochondrial translation assay

Cells were seeded in the DMEM medium in 10 cm^2^ 6-well plates pre-treated with 0.1% gelatine 24 h before the experiment. The cells were incubated in the RPMI medium without methionine (Gibco) containing 100 µg/ml emetine (Carl Roth) for 10 min in the cell incubator at 37°C and 5% CO_2_. Then 140 µCi/ml EasyTag ^35^S-*L*-Methionine (1,175 Ci/mmol, Perkin Elmer) was added and cells were incubated for another 30 min. After that 150 µg/ml cold methionine was added for 10 min. Then cells were collected with ice-cold 1× PBS, pelleted and resuspended in the same buffer. An aliquot was used for protein measurement, the rest was dissolved in Laemmli buffer and sonicated. Samples were heated at 40°C for 3 min and resolved by 12% SDS-PAGE. Then proteins were transferred onto an Amersham Protran western blotting nitrocellulose membrane (GE Healthcare) by semi-dry transfer. The membrane was dried and the radioactive signal was visualised with Phosphorimager on Typhoon Trio (GE Healthcare). Images were analysed with ImageQuant TL (v. 7.0, GE Healthcare). After this, western blots were performed as described above.

### Mitochondrial DNA copy number measurement

Cells were seeded on a 6-well plate in standard medium, harvested at subconfluency with 1× PBS and dissolved in PKLB buffer (50 mM Tris-HCl, pH 8, 1 mM EDTA, 0.5% Tween-20) supplemented with 100 µg/ml proteinase K and 100 µg/ml RNase A. After incubation for 2 h at 55°C with agitating at 1,100 rpm, the lysate was boiled for 10 min at 95°C and diluted 1:20 in 0.5× TE buffer, pH 8, to use directly in a probe-based real-time PCR. TaqMan probes specific for *MT-ND1* (mitochondrial DNA-encoded gene) and 18S rRNA (nuclear reference for normalization) were used together with primers 5mtDNA(ND1)1/3mtDNA(ND1)1 (36) and 5_18SrRNA1/ 3_18SrRNA1 (22), respectively.

### RT-qPCR

Cells were harvested and washed with 1× PBS. Pellets were lysed in RNAzol RT (Molecular Research Center) and total RNA was prepared according to the manufacturer’s instructions. For analysis, 250 ng of RNA were treated with DNase I (Thermo Fisher Scientific) and used for cDNA synthesis and RT-PCR, as described in (22,37). RNA levels were determined by real-time qPCR with primers specified in Table S3 and normalized to 18S rRNA and *UBC* mRNA levels.

### RNA stability measurements

Cells were seeded at subconfluency in 6-well plates, where they were exposed to 250 ng/µl ethidium bromide in standard medium to block mitochondrial transcription. For the stability assay, samples from treated cells were taken at 4, 6, 8, 10 and 24 hours and directly lysed in RNAzol for RNA preparation and kept at -20 °C. RNA was extracted and analysed by RT-qPCR, as described in the corresponding section. RNA levels determined by real-time qPCR were normalized to 18S rRNA and *UBC* mRNA and related to the time point 0.

### Transcriptomic analyses

Mitochondria of the three *YBEY*^+^ and three *YBEY* knockout clones were isolated as described above, dissolved in an equal volume of RiboZol (Amresco), and total RNA was prepared according to the manufacturer’s instructions. RNA was dissolved in 0.5× TE buffer, pH 7.5, and used for cDNA library generation with the NEBNext Ultra kit (New England BioLabs), without enrichment, with short fragmentation, the cutoff size 200-800 bp, followed by Illumina HiSeq V4 paired-end sequencing at the VBCF core facility in Vienna. Sequencing read libraries were converted from the BAM format to the FASTQ format with the bamtofastq tool from the BEDTools2 package (v2.26) (38), processed with cutadapt (version 1.15) (39) and then mapped to the *H. sapiens* mitochondrial genome (Genbank accession J01415.2) with the use of segemehl (version 0.1.7) (40) with -q and -p options for paired reads. The resulting alignment files in the SAM format were converted to BAM files using the Samtools subcommand view (version 1.4) (41) with the flags -bS. Feature quantification was done using featureCounts (version v1.4.6) (42) from the Subread package with the options -T (10) -p -t (exon) -0 -g (gene_id) -s 1 with a manually curated human mitochondrial genome annotation to generate raw read counts. Differential gene expression analysis was performed with the DESeq2 package (version 1.16.1) (43) in R version 3.4.4. RNA-seq tracks were visualized in the Integrated Genome Browser (v. 9.0.2) (44).

### cRT-PCR

Total RNA was isolated from 10 cm^2^ of cells grown standardly to 80-90% confluence with TRIzol reagent (Invitrogen), following the manufacturer’s protocol. To ensure that all processed transcripts have ligation-competent 5’-phosphate and 3’-hydroxyl termini, 5 µg of RNA was treated with 10 U of T4 polynucleotide kinase (Thermo Fisher Scientific) in the presence of 1 mM ATP and 50 U of RNaseOUT (Thermo Fisher Scientific) in 1× Buffer A (50 mM Tris-HCl, pH 7.5, 10 mM MgCl_2_, 5 mM DTT, 0.1 mM spermidine), according to the manufacturer’s protocol, followed by P:C:I extraction.

For circularisation, a 20 µl reaction, containing 300-1,000 ng of RNA, 50 U of RNaseOUT, and 10 U of T4 RNA ligase (Thermo Fisher Scientific) in 1× RNA ligase buffer (50 mM Tris-HCl, pH 7.5, 10 mM MgCl_2_, 10 mM DTT, 1 mM ATP), was assembled and incubated overnight at 16-18°C. In parallel, a control reaction without RNA ligase was performed. Both RNA samples were P:C:I-extracted and 100-400 ng of RNA was used as templates for RT-PCR.

RT-PCRs were performed with the One-step RT-PCR kit (QIAGEN) with SAO00187/SAO00188 (for 12S rRNA), SAO00204/SAO00205 (for 16S rRNA), or SAO00191/SAO00192 (for the *MT-CO2* mRNA) oligonucleotides as primers with the default manufacturer’s protocol. A portion of the RT-PCR samples were visualized by agarose gel electrophoresis followed by ethidium bromide staining to ensure that only circularized RNA gave rise to amplification products. The rest was purified with the GeneJET PCR purification kit (Thermo Fisher Scientific), digested with XbaI/KpnI (for 12S and 16S rRNAs) or PstI/KpnI (for the *MT-CO2* mRNA), and P:C:I-extracted.

For cloning, 150-300 ng of the digested RT-PCR product was mixed with 30-50 ng of pUC19, digested with the same restriction endonucleases and dephosphorylated with FastAP (Thermo Fisher Scientific), in the presence of 1 Weiss U of T4 DNA ligase (Thermo Fisher Scientific) in 1× Ligation buffer (40 mM Tris-HCl, pH 7.8, 10 mM MgCl_2_, 10 mM DTT, 0.5 mM ATP) and incubated on bench overnight. The resulting ligation mixes were used to transform competent XL1 Blue cells. pUC19-containing clones were selected on solid LB medium with 100 µg/ml ampicillin. Typically, ∼50 colonies per plate were screened for the presence of the insert by colony PCR with the M13 reverse and M13 (−21) forward primers. Plasmids from positive clones were sequenced. Sequences were analysed with Chromas (v. 2.6.2) and aligned versus the reference *H. sapiens* mitochondrial genome (Genbank accession J01415.2) with BLAST.

### Coimmunoprecipitation assay

For the experiment shown on Figure 5A and Table S6, SAL001 cells inducibly overexpressing YBEY-3×FLAG were used, while the parental Flp-In T-REx 293 cell line served as negative control. In both cases, 900 cm^2^ of cells were grown and induced, as described in section “Human cell lines”. Cells were harvested with ice-cold 1× PBS, and crude mitochondria were isolated as above and lysed in 0.5 ml of 20 mM Tris-HCl, pH 7.5, 150 mM KCl, 1 mM MgCl_2_, 1 mM CaCl_2_, 1 mM DTT, 0.5% n-dodecyl-β-D-maltoside, 1 mM PMSF with the help of a Dounce homogeniser (20 strokes). This and the subsequent manipulations were carried out at 4°C. The lysate was cleared at 14,000g for 10 min and mixed with 50 µl of the α-FLAG antibody (Sigma-Aldrich, Cat#F1804), followed by incubation for 30 min with continuous rocking. Thereafter, the lysate was mixed with 100 µl of protein A-sepharose beads (Sigma-Aldrich), pre-washed five times with 1 ml of 20 mM Tris-HCl, pH 7.5, 150 mM KCl, 1 mM MgCl_2_, 1 mM CaCl_2_, 1 mM DTT, 1 mM PMSF in a 1.5 ml tube, and incubated for another 30 min with continuous rocking. The beads were collected at the bottom of the tube by pulse centrifugation, and the flow-through was discarded. The beads were washed five times with 1 ml of 20 mM Tris-HCl, pH 7.5, 150 mM KCl, 1 mM MgCl_2_, 1 mM CaCl_2_, 1 mM DTT, 1 mM PMSF, and the retained proteins were eluted with 150 µl of 1× Laemmli buffer by boiling at 80°C for 5 min. For western blot analysis, 40 µl of the eluates in parallel with the corresponding cleared lysate, flow-through and wash fractions (equivalent of 1/500 of the starting material) were resolved by 15% SDS-PAGE followed by western blotting as described above.

For LC-MS/MS analysis, the eluted proteins were precipitated overnight with five volumes of glacial 0.1 M ammonium acetate in 100% methanol. After centrifugation at 12,000g at 4°C during 15 min, the resulting pellets were washed twice with 0.1 M ammonium acetate in 80% methanol and dried under vacuum (Genevac centrifugation concentrator miVac Duo, Fisher Scientific). The pellets were resuspended in 100 µl of 50 mM ammonium bicarbonate and submitted to reduction (5 mM dithiothreitol, 95°C, 10 min) and alkylation (10 mM iodoacetamide, room temperature, 20 min). Proteins were finally digested overnight with 200 ng of sequencing grade trypsin (Promega). Samples were analyzed with nanoLC-MS/MS on either a TripleTOF 5600 mass spectrometer (SCIEX) coupled to a NanoLC-Ultra-2D-Plus system (Eksigent), as described previously (45) or on a Q-Exactive Plus mass spectrometer (Thermo Fisher Scientific) coupled to an EASY-nanoLC-1000 chromatograph (Thermo Fisher Scientific) as described previously (46). Each sample was separated on an analytical Acclaim PepMap 100 C18 LC column, 3 µm, 250 mm length, 0.075 mm I.D. (Thermo Fisher Scientific) with a 160 min 300 nL/min gradient of acetonitrile. MS data were searched against the *Homo sapiens* Swissprot database with a target-decoy strategy (Swissprot release 2019_07; 20432 protein sequences). Peptides and proteins were identified with the Mascot algorithm (Matrix Science) and data were further imported into Proline (ProFI Proteomics, http://proline.profiproteomics.fr/). Proteins were validated on Mascot pretty rank equal to 1, and 1% FDR on both peptide spectrum matches (PSM score) and protein sets (Protein Set score). Proline was used to align the spectral count values across all samples using the “Compare with SC” tool. The total number of MS/MS fragmentation spectra was used to quantify each protein.

Overall, 5,233 peptides (from 708 proteins), 3,802 peptides (from 532 proteins), 13,620 peptides (from 895 proteins), and 4,353 peptides (from 567 proteins) were identified in the control #1, control #2, YBEY-3×FLAG coIP #1, and YBEY-3×FLAG #2 samples, respectively. To establish the list of YBEY-3×FLAG-associated candidate proteins, highly variable and ambiguous proteins were first filtered. Then the number of spectral counts for each protein was normalized by the total number of spectral counts in the sample, and ratios YBEY-3×FLAG coIP / control coIP were taken. Proteins enriched above two medians of these ratios were retained. Poisson 99% confidence intervals for YBEY-3×FLAG coIP spectral counts were built with the help of GraphPad QuickCalcs, and the proteins for which the 99% CI did not include the negative control value were retained. The list of proteins reproducibly enriched in both experiments according to these criteria was further filtered with additional YBEY coIPs to remove likely false positives. The remaining proteins were declared candidate YBEY binders (Table S6).

Spectral abundance factors (SAF) for YBEY and p32 were calculated as described in (47) by normalising the number of the spectral counts by the length of the mature protein (in amino acids). The ratio between the two SAFs provides an estimate for the YBEY:p32 stoichiometry.

The experiment shown on Supplementary Figure 8B was carried out largely in the same way, with the following alterations. As biological material, 1,800 cm^2^ of Flp-In T-REx 293 cells were used to isolate crude mitochondria. The mitochondria were lysed as above in 1 ml of 20 mM Tris-HCl, pH 7.5, 150 mM KCl, 1 mM MgCl_2_, 1 mM CaCl_2_, 1 mM DTT, 0.5% n-dodecyl-β-D-maltoside, 1 mM PMSF, and the cleared lysate was split in two 0.5 ml portions. One portion was mixed with 50 µl of the α-p32 antibody (Santa Cruz Biotechnology, Cat#sc-271200), whereas the other was treated with 50 µl of the α-FLAG antibody, which in this case served as negative control.

The experiment shown on Supplementary Figure 8A was performed as follows. We first established four parallel cell cultures of 225 cm^2^ each. The first one was the parental Flp-In T-REx 293 cell line transfected with the empty pcDNA5 FRT/TO plasmid, the second – the same cell line transfected with pSAP0033 to overexpress p32-HA, the third – the SAL001 cell line transfected with the empty pcDNA5 FRT/TO plasmid to inducibly overexpress YBEY-3×FLAG, the fourth – the same cell line transfected with pSAP0033 to have both YBEY-3×FLAG and p32-HA overexpressed. (The pSAP0033 plasmid was obtained by cloning the *C1QBP* cDNA, amplified with primers SAO00096/SAO00097, into the BspTI/XhoI sites on pcDNA5 FRT/TO.) These cells were grown to 70-80% confluency and transfected with 22.5 µg of the corresponding plasmid (mixed with 60 µl of Lipofectamine 2000 in 20 ml OptiMEM containing 0.2 µg/ml tetracycline) for 6 h. The medium was then changed to EMEM and the cells were harvested in 18 h with ice-cold PBS and directly lysed in 0.5 ml of 20 mM Tris-HCl, pH 7.5, 150 mM KCl, 1 mM MgCl_2_, 1 mM CaCl_2_, 1 mM DTT, 0.5% n-dodecyl-β-D-maltoside, 1 mM PMSF. The resulting cleared lysates were split in two portions of 250 µl, each mixed with either 10 µl of the α-FLAG antibody or 10 µl of the α-HA antibody (Sigma-Aldrich, Cat#H6908) and incubated for 30 min with constant rocking. Thereafter, the lysates were mixed with 40 µl of protein A-sepharose beads, pre-washed as above. The rest of the procedure was the same. The cleared lysate, flow-through, wash fractions (equivalent of 1/125 of the starting material) and eluates (20 µl) were analysed by 15% SDS-PAGE followed by western blotting.

### Purification of proteins from *E. coli*

*YBEY, C1QBP, MRPS11* and *MRPL18* cDNAs were obtained by RT-PCR from total RNA (isolated from Flp-In T-REx 293 cells, as described in section “cRT-PCR”). RT-PCRs were performed with the One-step RT-PCR kit by the default manufacturer’s protocol with the following primers: SAO00143/SAO00144 (*YBEY*), SAO00173/SAO00174 (*C1QBP*), SAO00213/SAO0213 (*MRPS11*), SAO00161/SAO00163 (*MRPL18*). The resulting RT-PCR products were digested with corresponding restriction endoribonucleases and used for cloning as follows. *YBEY* was first cloned into pET-21d via the NcoI and XhoI sites with a C-terminal His_6_ tag, and the resulting inhibitory short upstream ORF was removed by QuikChange with SAO00153/SAO00154 to derepress the YBEY protein production. The resulting pSAP0077 plasmid conferring resistance to ampicillin was introduced into Rosetta cells to yield the IPTG-inducible SAB0103 strain. The same plasmid served as template to derive YBEY mutant versions by QuikChange with the following primers: SAO00042/SAO00043 (R55A), SAO00056/SAO00057 (R57A), SAO00044/SAO00045 (D62A), SAO00046/SAO00047 (H118A), SAO00048/SAO00049 (H122A), SAO00050/SAO00051 (H128A), SAO00052/SAO00053 (M137A), SAO00054/SAO00055 (E141A). The resulting plasmids (pSAP0079 through pSAP0086, respectively) were introduced in Rosetta cells to yield the SAB0116 through SAB0123 strains.

*C1QBP* (mature form) was cloned into pET-30a via the NdeI and XhoI sites without any tag. The resulting pSAP0097 plasmid conferring resistance to kanamycin was introduced into Rosetta cells to yield the IPTG-inducible SAB0144 strain. The SAB0146 strain, containing both pSAP0077 and pSAP0097 and, therefore, inducibly co-overexpressing both YBEY-His_6_ and p32, was constructed by transforming SAB0144 with pSAP0077.

*MRPS11* (mature form) was first cloned into pET-30a via the NdeI and XhoI sites with a C-terminal His_6_ tag, which was removed by QuikChange with SAO00221/SAO00222 primers. The resulting pSAP0112 plasmid conferring resistance to kanamycin was introduced into Rosetta cells to yield the IPTG-inducible SAB0199 strain.

*MRPL18* (mature form) was cloned into pET-15b via the NdeI and BamHI sites with an N-terminal His_6_ tag, and the ORF was corrected by QuikChange with SAO00064/SAO00065 primers. The resulting pSAP0007 plasmid conferring resistance to ampicillin was introduced into BL21 Star (DE3) cells to yield the IPTG-inducible SAB0021 strain. The identities of all inserts were verified by Sanger sequencing.

To purify YBEY-His_6_ variants or His_6_-uL18m under denaturing conditions, the SAB0103/ SAB0116-SAB0123 or SAB0021 strains, respectively, were grown standardly overnight in the LB medium at 37°C in the presence of 100 µg/ml ampicillin and 34 µg/ml chloramphenicol. The resulting culture was inoculated 1:100 in 10 ml of fresh LB medium with the same antibiotics and grown at 37°C at constant shaking until OD_600_ of 0.5-0.8. Then 0.1-0.2 mM IPTG was added to induce protein production, the culture was shifted to 30°C and grown for another 4 h. The induced culture was rapidly cooled down on ice for 10 min and harvested by centrifugation at 4,000*g* for 10 min at 4°C. The cell pellet was washed thrice with ice-cold 50 mM Tris-HCl, pH 8, 150 mM KCl. The pellet was resuspended in 500 µl of denaturing lysis buffer (50 mM Tris-HCl, pH 8, 500 mM KCl, 8 M urea) at room temperature and sonicated until the solution became clear. The lysate was cleared by centrifugation at 14,000*g* for 10 min at room temperature. The cleared lysate was loaded on 100 µl of Ni-NTA agarose beads (QIAGEN), pre-washed twice with 1 ml of denaturing washing buffer (50 mM Tris-HCl, pH 7.5, 500 mM KCl, 1 mM MgCl_2_, 20 mM imidazole, 8 M urea), and incubated with constant rocking for 1-2 h at room temperature. The beads were sedimented by pulse centrifugation and the flow-through was discarded. The beads were washed twice with 1 ml of denaturing washing buffer at room temperature. Snap renaturation was performed by triple washing with 1 ml of native washing buffer (20 mM Tris-HCl, pH 7.5, 150 mM KCl, 1 mM MgCl_2_, 40 mM imidazole, 1 M urea, 0.05% Triton X-100, 1 mM PMSF) at 4°C. The residual urea in this and the subsequent buffers, insufficient to denature proteins, was found to improve YBEY solubility. Elution was achieved by step-wise addition and removal of 250 µl of elution buffers containing 20 mM Tris-HCl, pH 7.5, 150 mM KCl, 1 mM MgCl_2_, 1 M urea, 0.05% Triton X-100, 1 mM PMSF, and 50 mM, 60 mM, 70 mM, 80 mM, 100 mM, 120 mM, 140 mM, 160 mM, or 180 mM imidazole at 4°C (5 min per step). Low concentrations of imidazole preferentially remove nonspecifically bound proteins, whereas YBEY-His_6_ and His_6_-uL18m are eluted mostly at 140-160 mM imidazole. These two fractions were typically used for enzymatic assays (see section “*In vitro* RNase assays”). The purity of purified proteins was confirmed by SDS-PAGE and LC-MS/MS. The purified proteins were used immediately or stored at -80°C for no more than 1 month (longer storage often results in YBEY-His_6_ aggregation and inactivation).

For protein purification under native conditions from the SAB0103, SAB0144, SAB0146 and SAB0199 strains, 40 ml cultures were grown and induced as above. The pellet of harvested bacteria was resuspended in 500 µl of ice-cold native lysis buffer (20 mM Tris-HCl, pH 7.5, 150 mM KCl, 1 mM MgCl_2_, 1 mM PMSF) and sonicated. The lysate was cleared as above. In the case of the ternary YBEY-His_6_-p32-uS11m complex reconstitution, the cleared lysates of the SAB0146 and SAB0199 bacteria were mixed 1:1 and incubated for 30 min at 4°C at constant rocking. The cleared lysate was loaded on 100 µl of Ni-NTA agarose beads, pre-washed twice with 1 ml of washing buffer (20 mM Tris-HCl, pH 7.5, 150 mM KCl, 1 mM MgCl_2_, 40 mM imidazole, 1 mM PMSF), and incubated at constant rocking for 1-2 h at 4°C. The beads were sedimented by pulse centrifugation and washed thrice with 1 ml of the ice-cold washing buffer. The proteins were eluted step-wise by addition and removal of 250 µl of elution buffers containing 20 mM Tris-HCl, pH 7.5, 150 mM KCl, 1 mM MgCl_2_, 1 mM PMSF, and 50 mM, 60 mM, 70 mM, 80 mM, 100 mM, 120 mM, 140 mM, 160 mM, or 180 mM imidazole at 4°C (5 min per step). Protein concentration was measured by Bradford assay.

### *In vitro* RNase assays

The templates for the 12S rRNA-mt-tRNA^Val^ and tRNA^Gln^ precursor substrates were obtained by PCR with primers SAO00085/HmT7tRNA_Val_R and SAO00149/SAO00150. The corresponding RNAs were produced with the T7 RiboMAX Express large scale RNA production system (Promega) and purified by denaturing PAGE. To perform the cleavage assay, ∼50 ng of RNA per sample were diluted in 10 µl of the reaction buffer (50 mM Tris-HCl, pH 7.5, 150 mM KCl, 1 mM MgCl_2_) and rapidly mixed with 5 µl of YBEY or uL18m solutions, diluted with the corresponding storage buffer (see section “Purification of proteins from *E. coli*”) to achieve the desired final concentration. The reactions were incubated at 25°C for 30 min (Supplementary Figures 6A and 7A). Alternatively, 15 µl samples were systematically retrieved from an accordingly scaled up reaction mixture after indicated times (Supplementary Figure 6B). Then 15 µl of denaturing RNA loading buffer (0.025% SDS, 18 mM EDTA, 0.025% bromophenol blue, 0.025% xylene cyanol in deionised formamide) was immediately added to each sample, and the reaction was stopped by boiling at 95°C for 5 min, followed by denaturing 6-8% 8 M urea PAGE in 1× TBE and northern blotting with HmT7tRNA_Val_R, SAO00093, or SAO00160 as probes.

### High-resolution glycerol gradient analysis

High-resolution gradient centrifugation was performed as in (48), with modifications to increase resolution, as previously suggested (49). For one gradient, 75 cm^2^ of cells grown standardly to 80-90% confluence were harvested and washed with ice-cold 1× PBS. The cell pellet was resuspended in 0.5 ml of the lysis buffer (50 mM Tris-HCl, pH 7.5, 150 mM KCl, 20 mM MgCl_2_, 0.5% β-dodecylmaltoside, 1 mM DTT, 1 mM PMSF) and lysed with the help of a Dounce homogeniser (20 strokes). This and the subsequent manipulations were carried out at 4°C. The lysate was cleared at 14,000*g* for 10 min and loaded on a linear 10-40% (w/v) glycerol gradient formed in SW32.1Ti ultracentrifugation tubes (Beckman Coulter) from the light (50 mM Tris-HCl, pH 7.5, 150 mM KCl, 20 mM MgCl_2_, 0.025% β-dodecylmaltoside, 10% glycerol (w/v), 1 mM DTT, 1 mM PMSF) and heavy (50 mM Tris-HCl, pH 7.5, 150 mM KCl, 20 mM MgCl_2_, 0.025% β-dodecylmaltoside, 40% glycerol (w/v), 1 mM DTT, 1 mM PMSF) solutions with the help of the Gradient Master 108 (Biocomp). The gradients were centrifuged for 17 h on an Optima XPN-100 ultracentrifuge (Beckman Coulter) at 26,000 rpm and fractionated in 30×550 µl fractions. For protein analysis, 90 µl of each fraction was mixed with 30 µl of 5× Laemmli buffer. The rest was treated with P:C:I and RNA was re-extracted with TRIzol. Proteins were analysed by western blotting. RNA was resolved by denaturing 6% PAGE and analysed by northern blotting.

### LC-MS/MS analysis of mitoribosomes

Mitochondria were prepared as described in section “Subcellular and submitochondrial fractionation”, with some modifications. 293T-REx cells (three technical replicates) were harvested from 600 cm^2^ and the three *YBEY* knockout clones from 1,200 cm^2^. Cell pellet was resuspended in 4 ml of chilled RSB buffer and incubated for 15 min on ice. Cells were homogenized with a small glass douncer. An equal volume of chilled MS buffer was added and the suspension was gently mixed. Nuclei and cell debris were removed and mitochondria pelleted as described above. The mitochondrial pellet was resuspended in 1 ml RSB/MS buffers (1:1), pelleted again, resuspended in 200 µl of M1 buffer (600 mM sucrose, 50 mM Tris-HCl, pH 7.6, 1 mM EDTA) and loaded on a sucrose step gradient (1 M sucrose over 1.5 M sucrose in 10 mM Tris-HCl, pH 7.6, 1 mM EDTA) formed in a 14×89 mm tube (Beckman Coulter). After centrifugation at 20,000*g* for 30 min at 4°C on a SW 41 Ti rotor (Beckman Coulter), the mitochondrial interphase was transferred to a fresh tube and an equal amount of 1× TE buffer was added. Mitochondria were pelleted at 10,000*g* at 4°C for 6 min and washed once with chilled M3 buffer supplemented with 1 mM DTT and 0.1% proteinase inhibitor cocktail. The mitochondrial pellet was resuspended in chilled M3 buffer supplemented with 1 mM DTT, 0.1% proteinase inhibitor cocktail and 0.02% digitonin and incubated for 10 min on ice. The mitochondrial pellet was washed once with chilled M3 buffer supplemented with 1 mM DTT and 0.1% proteinase inhibitor cocktail and stored at -80°C for further use.

The mitochondrial pellet was dissolved in 200 µl of lysis buffer (25 mM HEPES_KOH, pH 7.4, 100 mM KCl, 25 mM MgCl_2_, 2 mM DTT, 0.01% proteinase inhibitor cocktail and 1.7% Triton X-100) and incubated on ice for 15 min. After clearing the lysate by centrifugation at 30,000*g* for 20 min at 4°C, the lysate was loaded on a sucrose cushion (20 mM HEPES-KOH, pH 7.4, 100 mM KCl, 20 mM MgCl_2_, 2 mM DTT, 0.01% proteinase inhibitor cocktail, 1% Triton X-100, 1 M sucrose) in a 11×34 mm polypropylene tube (Beckman Coulter) and centrifuged for 6 hours at 55,000 rpm at 4°C on a TLS-55 rotor (Beckman Coulter) in a TL-100 ultracentrifuge (Beckman Coulter). The pellet was dissolved in RapiGest SF (Waters) and the protein concentration was determined (Bio-Rad protein assay, Bio-Rad). The mitoribosomal lysates were frozen in liquid nitrogen and analyzed at the Proteomics core facility of the Medical University of Vienna. Sample preparation and measurement were carried out as previously described (50). Briefly, peptides were separated on a C18 µ-Pillar-Arrayed-Column (PharmaFluidics) using the nanoRSLC UltiMate 3000 HPLC system (Thermo Fischer Scientific). The pillars had an interpillar distance of 2.5 µm and the separation was performed at a flow rate of 600 nl/min over a total separation path of 2 m using a 10 min isocratic step (5% of the following solution: 50% acetonitryl, 30% methanol, 10% 2,2,2-trifluoroethanol, 0.1% formic acid), followed by increasing amounts of the above-mentioned solution to 20% until 30 min, and 40% from 30 min to 60 min. Columns were flushed with 90% of the solution for 5 min until 65 min, followed by a column equilibration of 11 min. The measurement was repeated 3 times for each sample. Blank samples (injection of loading solvent) were run between sample injections for cleaning of the separation system and preventing carry-over. Mass spectrometric detection and MS/MS analysis was performed using the Q-Exactive Orbitrap BioPharma (Thermo Fisher Scientific). Electrospray ionization was achieved by using a voltage of 2 kV.

Database search was performed using Proteome Discoverer 2.2 (Thermo Fisher Scientific) with the Swissprot Human Database (version Jan. 2019). Proteins were detected with a Q-Exactive Plus Biopharma mass spectrometer and the data were analysed with Scaffold (v. 4.6.5, Proteome Software). Two missed tryptic cleavages were allowed and the false discovery rate was 1%. For statistical analysis, 2 or more peptides with individual assignments at 95% confidence were required.

### Protein sequence and structure analysis

Sequences of the canonical isoforms of the cloned proteins were retrieved from UniProt (https://www.uniprot.org/) and their corresponding cDNA sequences from NCBI Nucleotide (https://www.ncbi.nlm.nih.gov/nucleotide). Predictions of mitochondrial targeting were performed with MitoFates (51), TargetP (52), and Mitoprot (53) with default parameters. Mature forms of YBEY, uL18m and p32 were chosen based on published data (19,54,55). The mature form of uS11m was arbitrarily chosen based on MS data retrieved from neXtProt (https://www.uniprot.org/), further refined with the MTS prediction tools and multiple sequence alignment of bacterial and mitochondrial uS11 orthologues, which were retrieved from InterPro (https://www.ebi.ac.uk/interpro/beta/), with COBALT (56). Structural model of human YBEY was obtained with RaptorX (57) with default parameters.

All protein and ribosome structures (58,59) were visualised in PyMOL (https://pymol.org/2/). Positionally equivalent YBEY residues for mutagenesis were identified by multiple sequence alignment (COBALT) of YBEY homologues retrieved from InterPro, further confirmed by the RaptorX structural model analysis with PyMOL. pI predictions were performed with the ExPASy: Compute pI/Mw tool (60).

### Phylogenetic analyses

Phyla with at least one genome sequenced with contigs of >100,000 bp (as retrieved from NCBI Genome, https://www.ncbi.nlm.nih.gov/genome, in January 2019) were visualised with the help of the Interactive Tree of Life (61). YBEY protein sequences were retrieved from InterPro and verified for contamination with NCBI BLAST (62). For apparent YBEY-negative groups, targeted searches were performed in NCBI BLAST with *E. coli*, human or the closest relative YBEY orthologue as queries. Obtained new hits were verified for the presence of the diagnostic features of the YBEY family (e.g. histidine triad).

### Statistical analyses

Full details of statistical measures, sample sizes, definitions and analyses are provided in figure legends or directly in the corresponding figures. Most of the descriptive statistics in this study, as well as Pearson’s correlation measurements, were done in Microsoft Excel 2010 (Microsoft), with the exception of the violin plots on Supplementary Figure 11 created with Microsoft Power BI Desktop (v. 2.71.5523.821). Two-tailed Fisher’s exact test was calculated in GraphPad QuickCalcs (GraphPad). Kolmogorov-Smirnov test was calculated in Physics: Tools for science (College of St Benedict, St John’s University; http://www.physics.csbsju.edu/stats/KS-test.n.plot_form.html). In the majority of cases, when *n* was high, the two-tailed Mann-Whittney test, free of the assumptions of normality and homoscedasticity, was used to compare groups. For low-to-moderate *n*, when the assumption of normality was met (i.e. qPCR-based data), the two-tailed unpaired Welch’s test, free of the assumption of homoscedasticity, was used to compare two means. In few remaining cases with *n* = 3 and homoscedastic normal data, for the sake of higher power, the one- or two-sample unpaired two-tailed t-test was applied. All these tests and the corresponding assumptions were implemented with the help of Statistics Calculators (Statistics Kingdom; http://www.statskingdom.com/index.html). Power and sample size calculations were performed in Power and Sample Size (HyLown Consulting LLC; http://powerandsamplesize.com/). Adjustments for multiple comparisons following the Benjamini-Hochberg or Bonferroni procedures were performed as described in (63).

## RESULTS

### Human YBEY is a mitochondrial protein

Controversy exists with regard to the subcellular localisation of human YBEY. One high-throughput study reported it as a nuclear (64) and another one as a mitochondrial protein (20). However, algorithms like MitoFates, TargetP and Mitoprot consistently predict its mitochondrial localization as well as an N-terminal mitochondrial targeting sequence (Figure 1A) (51,53,65). Indeed, subcellular fractionation revealed the presence of the protein in the mitochondrial but not in the nuclear fraction (Figure 1B). We also tagged the protein on its C-terminus with the FLAG epitope and tracked the protein by immunostaining in transiently or stably transfected human cells (Figure 1C, Supplementary Figure 2). The protein colocalized with mitochondria in all studied cell types, without evidence for an additional nuclear localization. Moreover, high-resolution confocal microscopy analysis suggested that YBEY localises in the inner compartment of mitochondria (Figure 1C,D). Upon submitochondrial fractionation with proteinase K treatment, YBEY behaved essentially like the matrix-localised mitoribosomal protein uL4m, showing complete resistance to digestion and only disappearing when the mitochondria were lysed (the outer membrane protein TOMM20, facing the cytosol, was digested in all analysed samples, whereas the inner membrane protein OPA1, which is highly exposed in the intermembrane space, specifically disappeared upon the rupture of the outer membrane; Figure 1E). Altogether, these experiments confirmed that YBEY is a *bona fide* mitochondrial matrix protein.

### YBEY is required for normal mitochondrial morphology and respiration

To study the cellular function of YBEY, we generated complete CRISPR knockouts of its gene (henceforth *YBEY* KO) in HEK293T-REx cells (Supplementary Figure 3). Compared to *YBEY*-expressing parent cells, *YBEY* KO cells showed a significant growth delay and fast medium acidification in glucose-containing medium and failed to thrive on galactose (requiring respiration), suggesting a mitochondrial phenotype (Figure 2A). To assess the effect of YBEY loss on mitochondria, we first analysed mitochondrial morphology by transmission electron microscopy (Figure 2B,C). Whereas *YBEY*^+^ cells had normally shaped mitochondria with multiple cristae, *YBEY* KO mitochondria were significantly enlarged and often misshaped, contained less cristae and presented a wide variety of morphological defects, including internal membrane structures and electron-dense inclusions. Since such morphological abnormalities are usually indicative of mitochondrial dysfunction, we assessed the respiration phenotype of *YBEY* KO cells and observed a dramatic decrease in basal and maximal respiration, as well as ATP production rates (Figure 2D), consistent with a pronounced complex I and IV deficiency (Figure 2E). Altogether, the *YBEY* KO resulted in severe mitochondrial dysfunction and the loss of respiration in human cells.

### YBEY is essential for mitochondrial gene expression

To establish the molecular basis of the observed respiration phenotype, we evaluated the levels of select subunits of respiratory complexes in *YBEY* KO cells by western blotting (Figure 3A). Whereas some of the nucleus-encoded subunits, like SDHA, remained unchanged, the mitochondrial DNA (mtDNA)-encoded COX2 protein was almost undetectable in *YBEY* KO clones. Similarly, NDUFB8, a nucleus-encoded complex I subunit which assembles at a mid-late stage and depends on the presence of the mtDNA-encoded ND5 protein (66), was depleted. In line with these observations, metabolic labelling of mitochondrial translation products revealed a nearly complete inability of *YBEY* KO mitochondria to synthesise mtDNA-encoded polypeptides (Figure 3B). Since mtDNA levels were unaffected by the YBEY loss (Supplementary Figure 4A), this general translational shutdown indicates a mitochondrial gene expression defect.

We first analysed the mitochondrial transcriptome of *YBEY* KO cells (Table S5). Similarly to what had been observed in bacteria (6,9,10,12), the YBEY loss in human cells was associated with pervasive changes in mitochondrial RNAs (Figure 3C). Whereas some noncoding antisense RNAs produced from the L-strand appeared upregulated, levels of five mRNAs encoded on the H-strand (including *MT-CO1* and *MT-CO2* specifying the COX1 and COX2 proteins, respectively) were significantly decreased in *YBEY* KO cells (Figure 3C, Supplementary Figure 4B). Their downregulation was further corroborated by RT-qPCR and single-molecule FISH (smFISH) (Figure 3D, Supplementary Figure 4C). Since the applied RNA-Seq protocol does not permit to robustly evaluate the levels of mature tRNAs, we measured the expression of select mitochondrial tRNA species of varying abundance and from different genomic contexts by northern blotting and smFISH, but did not find significant differences between the *YBEY* KO and parental cell lines (Supplementary Figure 4C-E). Altogether, these observations suggest that, while YBEY loss does not seem to globally affect mtDNA transcription, the expression levels of several transcripts have been perturbed. This implicates YBEY, directly or indirectly, in post-transcriptional gene expression control in mitochondria. On the other hand, these deregulations cannot be *per se* responsible for the severe general translation phenotype observed in *YBEY* KO mitochondria (Figure 3B).

### YBEY is required to maintain the steady-state level of 12S rRNA, but not for rRNA processing

Intriguingly, we observed that the level of 12S rRNA (encoded by the *RNR1* gene) was markedly decreased in *YBEY* KOs, whereas the level of 16S rRNA (encoded by *RNR2*) remained unchanged (Figures 3C,D and 4A, Supplementary Figure S4C,D). Moreover, 12S rRNA was significantly less stable in *YBEY* KO cells, as compared to *YBEY*^+^ (Figure 4B). This molecular phenotype suggests a defect on the mitoribosomal SSU side, reminiscent of what was previously observed in bacteria (3-8). However, as pointed out above, rRNAs in mammalian mitochondria, flanked by tRNA genes without any intervening sequences (Figure 4A), are supposed to require only mitochondrial RNase P and RNase Z for processing (21-23). To directly assess a possible involvement of YBEY in mitochondrial rRNA processing, we mapped the termini of 12S rRNA by RNA-circularisation RT-PCR, cloning and sequencing (cRT-PCR). Neither the 5’-nor the 3’-termini of 12S rRNA molecules were significantly altered in *YBEY* KOs (Figure 4C). In fact, the *YBEY*^+^ and *YBEY* KO cell lines contained virtually indistinguishable populations of correctly processed and polyadenylated 12S rRNA molecules (Figure 4D, Supplementary Figure 5A), excluding the involvement of human YBEY in mitochondrial SSU rRNA processing.

Similarly, RT-qPCR did not reveal an accumulation of unprocessed tRNA precursors in *YBEY* KO mitochondria (Supplementary Figure 5B), and cRT-PCR analysis of the *MT-CO2* mRNA termini did not uncover any significant defect in its processing or polyadenylation (Supplementary Figure 5C-E). The downregulation of select mRNAs observed in *YBEY* KO cells (Figure 3C,D) may in fact stem from a lack of translation, as previously demonstrated in *E. coli* (17). Interestingly, 16S rRNA, although similarly well processed and polyadenylated in both *YBEY*^+^ and *YBEY* KO cells, showed frequent irregular truncations, in particular in the domain I, in the latter background (Supplementary Figure 5F-H). Such “hidden nicks” may be a result of a subtle destabilisation at the level of the large ribosomal subunit (LSU), which finds further support in the analysis of the mitoribosome assembly described below.

### YBEY has a promiscuous endoribonuclease activity

The finding that *YBEY* KO does not result in detectable misprocessing of 12S rRNA (or any other mt-RNA studied) prompted us to evaluate its RNase activity. Recombinant human YBEY-His_6_, purified under denaturing conditions and renatured, showed robust endoribonuclease activity on the 3’-minor domain of 12S rRNA and tRNA^Val^, a substrate analogous to the proposed target of bacterial YbeY at the 3’-end of the SSU rRNA (Supplementary Figure 6A). However, the primary cleavage site in this substrate was completely off the expected position, truncating the 12S rRNA moiety instead of separating it from tRNA^Val^. Moreover, with increasing enzyme concentration the degradation of the substrate proceeded until it was reduced to short oligonucleotides. We observed a similar behaviour on a variety of other RNA substrates (see, for instance, Supplementary Figure 6B), suggesting that on its own YBEY does not site-specifically cleave RNAs but instead acts as a promiscuous degradative RNase.

To further dissect its RNase activity, we created a series of alanine replacement mutations targeting select conserved residues, some of which have been implicated in catalysis in bacterial YbeY homologues (Figure 1A) (5,12). These mutations affected the RNase activity of human YBEY *in vitro* to different extents, substitutions H128A and E141A in the active site being the most detrimental (Supplementary Figure 7A). We tested the importance of H128 and R55, the latter supposedly involved in RNA binding, for the function of YBEY in human mitochondria by complementing the *YBEY* KO with stably integrated copies of WT or mutant *YBEY* genes. Surprisingly, all of the tested YBEY variants rescued the 12S rRNA depletion and mitochondrial translation phenotypes, and restored cellular respiration (Supplementary Figure 7B-D). Altogether, these data suggest that, although human YBEY shows endoribonucleolytic activity, it is intrinsically indiscriminate and apparently dispensable for its main role in sustaining mitochondrial translation.

### YBEY interacts with p32 and a distinct set of mitoribosomal proteins

The destabilisation of 12S rRNA (Figure 4B) in the absence of any apparent RNA processing alterations suggested that rather than rRNA maturation, a later step in the biogenesis of the mitochondrial SSU may be impaired in *YBEY* KO cells. Therefore, we set out to investigate the protein interactome of human YBEY. To facilitate the detection and the pulldown of the protein, we used an engineered tetracycline-inducible YBEY-3×FLAG cell line (Supplementary Figure 2), which only minimally (3.2 ± 0.6-fold, mean ± SD) overexpresses the protein, ensuring that the observed interactions are physiologically relevant. Coimmunoprecipitation assays followed by LC-MS/MS identified a reproducible set of enriched YBEY-3×FLAG-binding proteins (Table S6), which were further confirmed by western blotting (Figure 5A).

One top-scoring YBEY partner was the putative RNA-binding protein p32/C1QBP, critically required for mitochondrial metabolism and associated with a variety of mitochondrial diseases and cancer (67-76). The interaction between the two proteins was confirmed in cross-pulldown assays with the alternate use of YBEY-3×FLAG and p32-HA proteins as bait and prey (Supplementary Figure 8A). Moreover, pulldown of endogenous p32 selectively enriched endogenous YBEY, excluding any possible effect of tagging or overexpression (Supplementary Figure 8B). Finally, fluorescence lifetime imaging-based Förster resonance energy transfer (FLIM-FRET) confirmed the interaction of the two proteins in intact cells (Figure 5B).

Notably, when YBEY was used as bait, it systematically copurified stoichiometric amounts of p32 (one-sample t-test *P* = 0.45 for the 1:1 stoichiometry; *P* = 0.034 and 0.043 for the 2:1 and 1:2 alternatives, respectively; Supplementary Figure 8C). In contrast, p32, which appears to be a much more abundant protein (20), copurified only small amounts of YBEY. This suggests that whereas YBEY is quantitatively involved in stoichiometric complexes with p32, most of the latter is not associated with YBEY. To further study this interaction, we decided to reconstitute it in a heterologous system of *E. coli* (Figure 5C, Supplementary Figure 9A). While human YBEY-His_6_ alone was insoluble and, therefore, could not be purified under native conditions, native p32, due to its negative charge (pI = 4), was fully soluble, but not retained by the Ni-agarose beads due to the lack of a His_6_ tag. However, when both proteins were co-expressed, a significant proportion of YBEY-His_6_ was solubilised and copurified together with p32 in stoichiometric amounts (Figure 5C, Supplementary Figure 9A), in line with our coimmunoprecipitation results. Altogether, these data suggest the existence of a stable and stoichiometric complex of the two proteins.

Most of the other high scoring proteins copurifying with YBEY were constituents of the mitochondrial ribosome (Figure 5A, Table S6). Their specific association with YBEY in intact cells was confirmed by FLIM-FRET analysis (Figure 5B); only one of these proteins, uS11m, is a component of the mitochondrial SSU, whereas the remaining nine LSU proteins all associate with or near domain I of 16S rRNA (Figure 5D). The interaction between YBEY and these mitoribosomal proteins appears to take place outside the mitoribosome, since endogenous YBEY formed small monodisperse complexes and did not associate with ribosomal particles upon glycerol gradient sedimentation (Supplementary Figure 9B). Moreover, it did not interact with other mitoribosomal proteins, such as mS35, mS37 and mL38 (Figure 5A,B). Given that the vast majority of the YBEY-associated LSU proteins are spatially clustered (Figure 5D) and recruited at a very early stage of the LSU assembly (77-79), this interaction may occur at the level of an early LSU assembly intermediate.

As pointed out above, the destabilisation of the 12S rRNA in YBEY KO suggested a defect on the side of the mitochondrial SSU. Thus, we decided to study the interaction between YBEY and uS11m in more detail. Recent studies in *E*.*coli* suggested that the function of YbeY may be tightly connected with two SSU-related proteins, uS11 and the ribosome biogenesis GTPase Era (80,81). Although the human Era homologue, ERAL1, was not coimmunoprecipitated with YBEY (Figure 5A), probably due to the GTP dependence of its interactions (82), both uS11m and ERAL1 were found to interact with YBEY *in situ* by FLIM-FRET (Figure 5B). Since no other SSU ribosomal proteins were recovered in our screen, we wondered whether YBEY interacts directly with uS11m outside the mitoribosomal context. To directly address this hypothesis, we reconstituted the interaction between recombinant YBEY-His_6_, p32 and uS11m produced in *E. coli* (Figures 5E, Supplementary Figure 9C). Strikingly, when lysates from bacteria coexpressing YBEY-His_6_ and p32 on the one side and uS11m on the other were coincubated, a stable and stoichiometric complex was formed. These results indicate that YBEY, p32 and uS11m strongly, directly and stoichiometrically interact with each other, raising the question of the functional significance of their association.

### YBEY loss results in uS11m-deficient mitoribosomes impaired in translation

To evaluate the effect of YBEY on its protein partners, we first analysed their abundance in *YBEY*^+^ and *YBEY* KO cells (Figure 6A). Whereas no significant changes were observed for p32, as well as several LSU proteins and some SSU proteins (mS27 and mS35), the mitoribosomal protein uS11m was strongly depleted in the *YBEY* KO. This finding was further corroborated by uS11m immunostaining in intact *YBEY*^+^ and *YBEY* KO cells (Figure 6B). Importantly, uS11m levels were restored by complementation of the *YBEY* KO with either WT or mutant *YBEY* genes (Figure 6C). Since the *MRPS11* mRNA level did not change significantly upon *YBEY* KO (Supplementary Figure S10), we hypothesised that uS11m may have been destabilised due to a deficiency in its incorporation into the mitoribosome, as it is frequently the case with other mitochondrial ribosomal proteins (78,83). Indeed, western blot analysis of high-resolution glycerol gradients revealed that, although the SSU in *YBEY* KO mitochondria sedimented similarly to its WT counterpart, it lacked uS11m, which could only be observed in small molecular weight fractions (Figure 6D). In line with this observation, the mitoribosomal protein mS37 was also strongly depleted in *YBEY* KO cells (Figure 6A). Together with bS21m, mS37 is an immediate neighbour of uS11m in the mitochondrial SSU (84,85), and whereas uS11m directly binds to 12S rRNA, the other two proteins structurally depend on uS11m (77), with which they form extensive protein-protein interactions (Figure 6E). Indeed, uS11m knockdown resulted in the same specific mS37 destabilisation as observed in *YBEY* KO cells (Figure 6F), confirming the hierarchical dependence of mS37 on uS11m.

**Figure 6:**
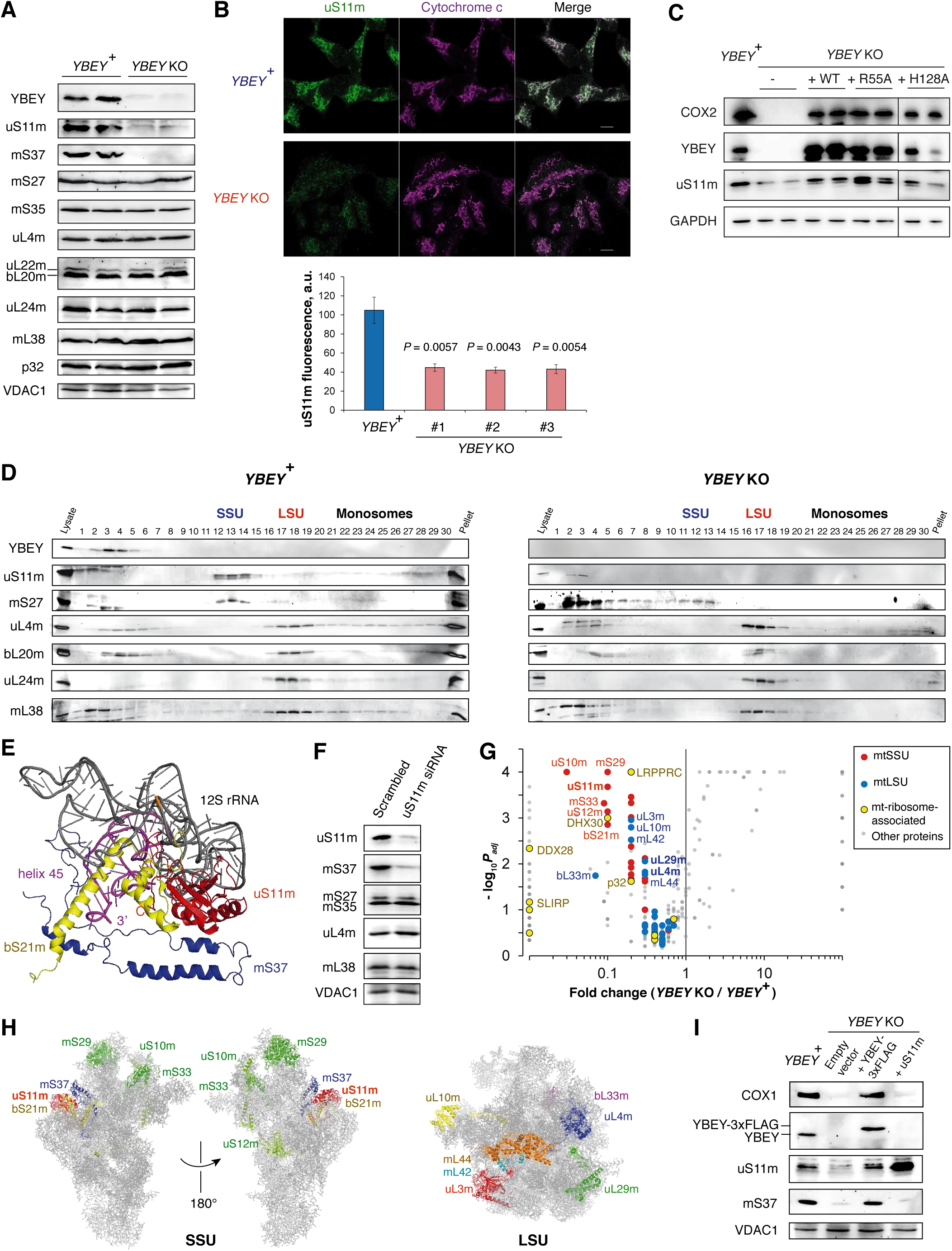
The YBEY loss results in uS11m-deficient mitoribosomes. **(A)** Western blot analysis of total protein from *YBEY*^+^ and *YBEY* KO cells reveals the selective depletion of mitoribosomal proteins uS11m and mS37. Two independent cell lines of each genotype are shown. VDAC1 is shown as loading control. **(B)** Quantitative immunofluorescence analysis of the uS11m protein levels in parental *YBEY*^+^ and three independent *YBEY* KO cell lines. Cytochrome c is shown as a mitochondrial marker. Means ± SEM for *n* = 6 (*YBEY*^+^ and *YBEY* KO #3), *n* = 5 (*YBEY* KO #1), and *n* = 7 (*YBEY* KO #2) frames are shown; *P*-values, two-tailed Welch’s test. **(C)** Rescue of uS11m and COX2 levels in *YBEY* KO cells by complementation with WT or mutant *YBEY* genes. Western blotting of total cellular protein is shown; GAPDH is used as loading control. All lanes are cropped from the same membrane. **(D)** High-resolution 10-40% glycerol gradient analysis of total *YBEY*^+^ and *YBEY* KO cell lysates followed by western blotting reveals the absence of uS11m from the mitochondrial SSU in *YBEY* KO cells. **(E)** The structural context of the uS11m protein in the mammalian mitochondrial SSU (pdb: 3jd5). The primary uS11m-binding site (residues 320-440 of 12S rRNA) is shown in grey. The helix 45 contacted by the C-terminal extension of uS11m (“C”) is shown in magenta. Proteins bS21m (yellow) and mS37 (blue) depend on extensive protein-protein contacts with uS11m and serve to sandwich the 3’-terminus of 12S rRNA (3’). **(F)** Transient knockdown of uS11m results in destabilisation of mS37. A non-targetting siRNA with a “scrambled” sequence was used as a control. **(G)** Quantitative mass spectrometry analysis of the levels of mitoribosomal and copurifying proteins from *YBEY* KO as compared to *YBEY*^+^ mitochondria. Particularly strongly downregulated proteins are named. See also Table S7. **(H)** Mapping of the proteins particularly strongly depleted from the mitoribosomes of *YBEY* KO cells on the structure of the mammalian mitoribosome. **(I)** Transient overexpression of YBEY-3×FLAG but not of uS11m in *YBEY* KO cells restores the translation of COX1 and rescues the stability of mS37. Western blotting of total cellular protein is shown.

To comprehensively evaluate the proteomic composition of the mitoribosomal subunits in *YBEY* KO cells, we purified them through a sucrose cushion and subjected them to quantitative LC-MS/MS (Figure 6G, Table S7). In line with the observed 12S rRNA depletion, levels of the mtSSU proteins were strongly decreased (two-tailed Mann-Whitney test *P* = 0.0013), whereas the overall abundance of mtLSU proteins was not significantly altered (*P* = 0.79). Looking at individual proteins, 7 LSU r-proteins were mildly but significantly depleted, suggesting a slightly perturbed LSU assembly (Figure 6H). This was paralleled by a decreased association of several LSU biogenesis factors, including DDX28 and DHX30 (86,87). In contrast, 6 SSU proteins were practically absent, including uS11m, bS21m and several other proteins mostly from the head of SSU (Figure 6H). These data confirm the SSU assembly defect in *YBEY* KO mitochondria suggested by our previous experiments and provide further support to the role of uS11m as a mechanistic link between YBEY and SSU biogenesis.

We wondered whether the depletion of cellular uS11m levels observed in *YBEY* KO cells was the direct cause of the SSU assembly defect. To this end, we replenished the uS11m pool by mildly overexpressing the protein from a plasmid in *YBEY* KO cells (Figure 6I). Strikingly, whereas re-expression of YBEY-3×FLAG in the same cells fully restored the uS11m and mS37 levels and COX1 production, uS11m overexpression failed to rescue mitochondrial translation and, importantly, did not restore the mS37 level. Together with the above results, this finding suggests that YBEY is not a mere uS11m stability factor, but is rather actively required for its incorporation in the SSU.

uS11m and its immediate neighbours bS21m and mS37 are key constituents of the mRNA exit channel (84,85). Therefore, a lack of these proteins would be expected to result in a translation initiation deficiency, explaining the observed translational shutdown in *YBEY* KO cells (Figure 3B). To assess this possibility, we analysed the association of mt-tRNA^Met^ and the *MT-ND3* mRNA, which form relatively small and clearly distinguishable extraribosomal RNPs, with the mtSSU in high-resolution glycerol gradients (Figure 7A). Whereas in *YBEY*^+^ cells a significant proportion of both RNAs cosedimented with the SSU, in *YBEY* KO cells neither RNA was found to be significantly SSU-bound (Figure 7A,B), suggesting that *YBEY* KO SSUs are indeed initiation-deficient.

**Figure 7:**
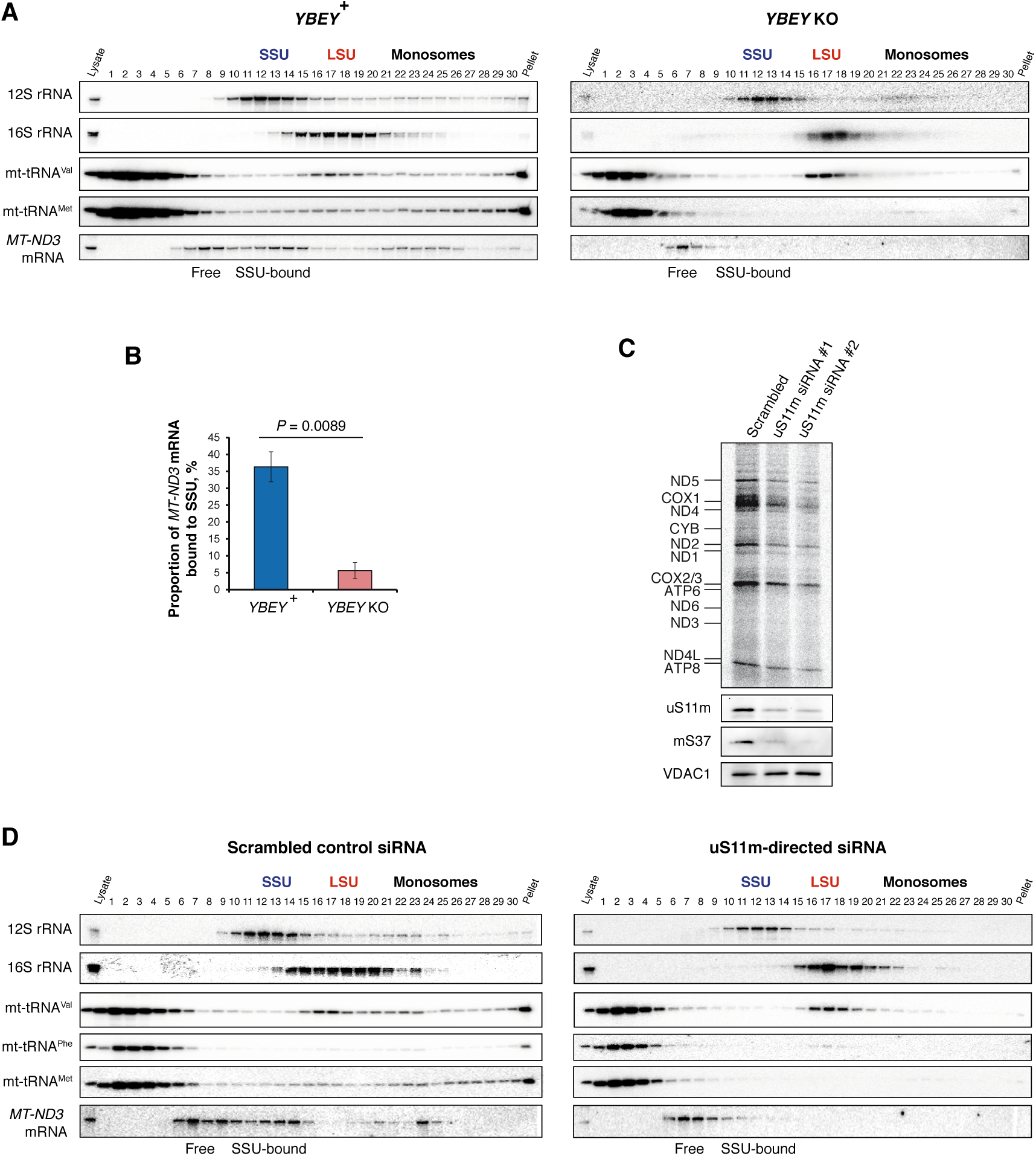
The knockdown of uS11m phenocopies the loss of YBEY. **(A)** High-resolution 10-40% glycerol gradient analysis of *YBEY*^+^ and *YBEY* KO total cell lysates followed by northern blotting shows a decreased association of the *MT-ND3* mRNA and mt-tRNA^Met^ with the mitochondrial SSU of *YBEY* KO cells. **(B)** Quantitative analysis of the association of extramitoribosomal *MT-ND3* mRNA with the SSU in the *YBEY* ^+^ and *YBEY* KO cells. Means ± SEM for *n* = 3 independent cell lines are shown; *P*-value, two-tailed Welch’s test. **(C)** Metabolic [^35^S]-methionine labelling of mitochondrial translation products reveals a strong decrease of protein synthesis upon transient uS11m knockdown. A non-targeting control siRNA (“Scrambled”) and two different uS11m-directed siRNA duplexes were used for HEK293T-REx cell transfection. The corresponding western blots confirm the uS11m knockdown and the concomitant depletion of mS37. VDAC1 is used as loading control. **(D)** High-resolution 10-40% glycerol gradient analysis of the total cell lysates from HEK293T-REx cells transiently transfected with control or uS11m-directed siRNAs followed by northern blotting shows a decreased association of the *MT-ND3* mRNA and mt-tRNA^Met^ with the mitochondrial SSU upon uS11m knockdown.

### uS11m knockdown phenocopies the *YBEY* KO mitochondrial translation phenotype

We reasoned that if *YBEY* KO effects are uS11m-mediated, uS11m knockdown should result in a similar translational phenotype. Indeed, transient RNAi-mediated uS11m depletion with two different siRNA duplexes significantly affected mitochondrial translation in HEK293T-REx cells and again resulted in the loss of mS37 (Figure 7C). Moreover, when we examined the association of mt-tRNA^Met^ and the *MT-ND3* mRNA with mtSSU in glycerol gradients, we acknowledged the same loss of mtSSU association as in *YBEY* KO cells (Figure 7D). Therefore, the uS11m knockdown largely phenocopies the *YBEY* knockout, further supporting the role of YBEY in recruiting uS11m in order to obtain initiation-competent mitochondrial SSUs.

## DISCUSSION

The deeply conserved YbeY protein has recently come into limelight as a critical factor required for normal physiology in Bacteria. Its loss has been associated with a wide array of debilitating phenotypes, such as sensitivity to abiotic stresses, inability to establish host-pathogen/symbiont relationships (6,7,13-15), metabolic deregulations and a severe growth impairment up to lethality (3,11,17). Widespread among Eukarya, YbeY has so far only been studied in *A. thaliana*, where it was found indispensable for chloroplast development and photosynthesis (16). Here we show that human YBEY localises to mitochondria and is essential for mitochondrial translation and, consequently, oxidative phosphorylation. We established that the inability of human cells to synthesise mitochondrial polypeptides in the absence of YBEY is apparently the consequence of a mitoribosome assembly defect: the severely destabilised mitochondrial SSU lacked uS11m and several other ribosomal proteins required for translation initiation. Based on interactome data and functional analyses, we propose that human YBEY, in complex with p32, acts to deliver uS11m to the nascent mitochondrial SSU in order to complete the assembly of an initiation-competent ribosomal particle (Figure 8).

**Figure 8:**
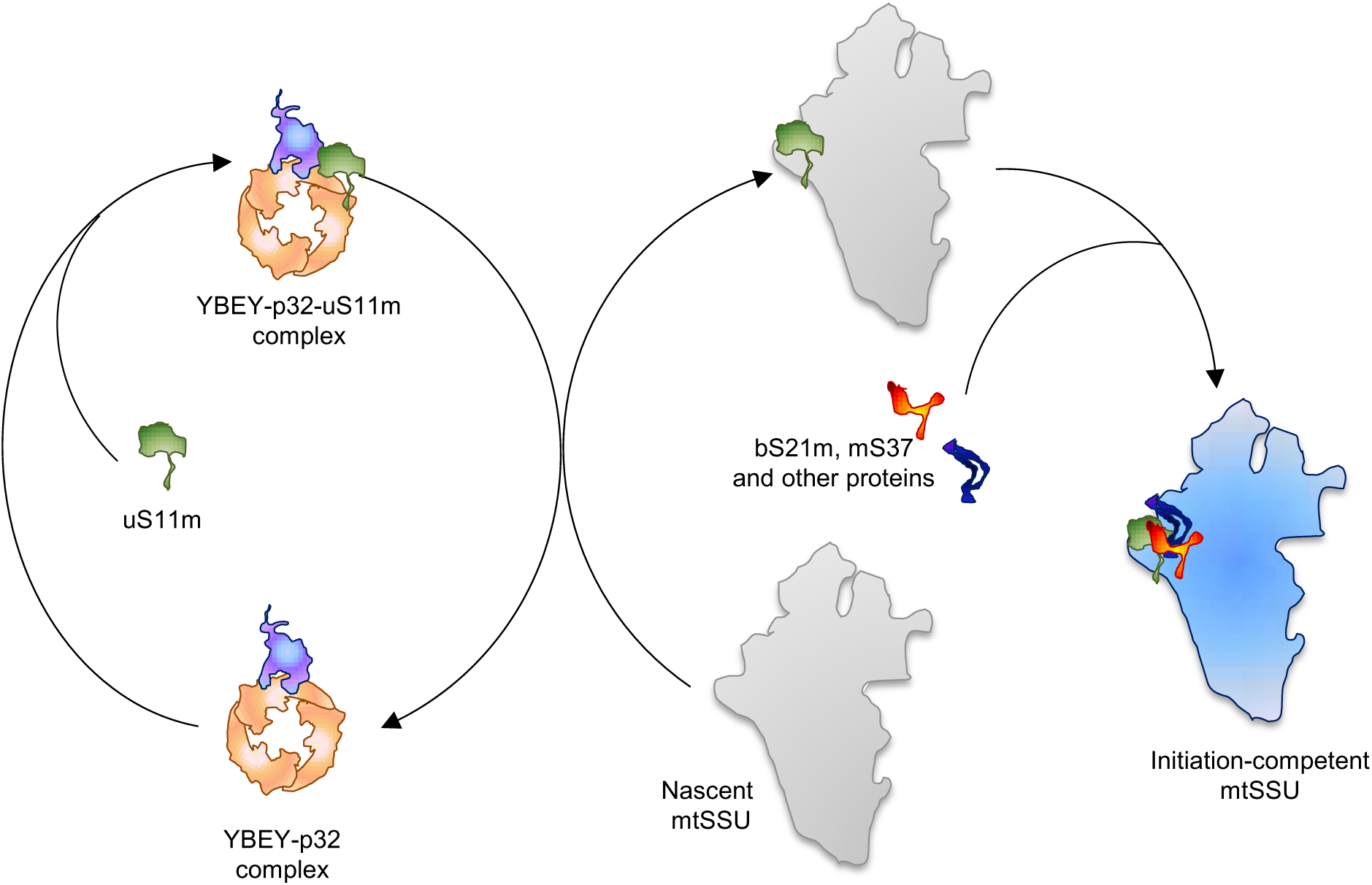
Human YBEY as an assembly factor for the mitochondrial SSU. For clarity, only one YBEY monomer per p32 trimer is shown, albeit our data support a heterohexamer model.

uS11m and its immediate neighbours on the mitochondrial SSU play key roles in translation initiation. uS11m, bS21m and mS37 line the mRNA exit channel (84,85). Additionally, uS11m makes extensive contacts with mtIF3 (88), predicting that the mtIF3 recruitment to mitoribosomes may be also perturbed in *YBEY* KO and uS11m knockdown cells. The bS21 protein is critically required for translation initiation in bacteria (89), and this function is likely maintained in mitochondria (84,85). Additionally, mS37 and bS21m are strategically positioned to lock the 3’-end of 12S rRNA (Figure 6E), which may contribute to its stability and the correct folding of the 3’-minor domain. All these features may explain why *YBEY* KO and uS11m knockdown cells seem to be deficient in mitochondrial translation initiation (Figure 7). Interestingly, the structural dependence of mS37 and bS21m incorporation on uS11m, associated with a considerable rearrangement of the SSU rRNA, has recently been demonstrated in *Trypanosoma bruicei* (90), suggesting that this assembly hierarchy is conserved in mitochondria across Eukarya. Given the deep conservation of YbeY, uS11, bS21 and Era and their interactions (80) in bacteria and bacteria-derived eukaryotic organelles, it will be important to verify whether the previously reported Δ*ybeY* translation phenotypes can be traced back to a similar SSU assembly defect, and especially the lack of uS11, as in human mitochondria.

*YBEY* KO mitochondrial SSU was found to lack several other functionally important proteins (Figure 6H) that are structurally connected with the uS11m-bS21m-mS37 module. mS29/DAP3 contacts mS37 via uS7m and plays a major structural role by forming the mitochondria-specific intersubunit bridges mB1a and mB1b (84). uS10m and mS33 form another structural module which is incorporated late during assembly and was proposed to hierarchically depend on the uS7/mS29-containing cluster (77). Interestingly, mS29 and uS10m were found to crosslink with mtIF3 (91), again suggesting that *YBEY* KO mitoribosomes may have a defect in mtIF3 recruitment and subunit association. While the studies of mitochondrial SSU assembly are still in their infancy (77,79,83), our data provide novel insights into the composition of a late SSU assembly intermediate and establish a hierarchical relationship between some of its constituents.

Interestingly, the molecular mechanism proposed here (Figure 8) shows striking parallels to the late stages of the cytosolic/archeal SSU biogenesis catalysed by the adenylate kinase Fap7/hCINAP (92). Similar to YBEY, Fap7 forms a stoichiometric complex with uS11/Rps14 (93) and helps to recruit it to the nascent SSU (94), which represents a final checkpoint before the fully assembled SSU is released for translation (95). In fact, all known uS11 homologues have a long disordered positively charged C-terminal extension, which needs to be correctly positioned in the vicinity of the helix-45 of the SSU rRNA (Figure 6E) (96). This requires the action of a dedicated chaperone (93). Fap7 also facilitates the association of the cytosolic uS11 with eS26 (97), which, by blocking the 3’-end of the 18S rRNA, is positionally and functionally analogous to the mitochondrial mS37 (84). Therefore, it appears that in human cells conceptually similar mechanisms operate to accomplish the assembly of both cytosolic and mitochondrial SSUs.

Several questions regarding the molecular mechanism of YBEY in human mitochondria remain to be answered. How is uS11m loading on the nascent SSU performed? What is the role of other YBEY-interacting factors, such as ERAL1 and p32? Involvement of ERAL1 in mitochondrial SSU biogenesis and the associated mitochondrial diseases were reported (98-100), and physical and genetic interactions between bacterial Era and YbeY were described (80,81). Moreover, the importance of Era for correct positioning of uS11 and recruitment of bS21 was demonstrated (82). Yet, the exact molecular role of ERAL1 in the YBEY pathway that we propose, is still elusive.

The major, stoichiometric partner of YBEY, p32, is a deeply conserved eukaryotic protein with a profound impact on the mitochondrial gene expression (67,75). Our purification experiments suggest that p32 enhances the intrinsically poor solubility of YBEY, which may be due to the unfavourable overall charge of human YBEY (pI = 7.04). Interestingly, most bacterial YbeY proteins, including those from Proteobacteria and Cyanobacteria, are highly negatively charged (Supplementary Figure 11). In contrast, eukaryotic YBEY homologues, with the exception of photosynthesizing clades, where YBEY appears to localise in chloroplasts (16), are neutral or slightly positive. Interestingly, this shift in charge coincides with the evolutionary emergence of the mitochondrial p32/MAM33 protein family (101), and it may help maintaining the solubility and negative charge of YBEY-containing complexes. The latter property is likely important for the interaction of YBEY with functionally relevant protein partners, such as uS11m.

The association of YBEY with LSU proteins (Figure 5A,B,D) and the truncations in domain I of 16S rRNA in *YBEY* KO cells (Supplementary Figure 5F,G), where these proteins bind, were further intriguing findings. Together with the decrease of some ribosomal proteins and LSU biogenesis factors in *YBEY* KO cells (Figure 6G,H), these observations suggest that LSU assembly is somewhat perturbed by the loss of YBEY too. Negative effects on the LSU side in the absence of YBEY have also been described in *E. coli* and in chloroplasts (8,16). It is, therefore, tempting to hypothesise the existence of a YBEY-mediated crosstalk between the SSU and LSU biogenesis pathways. Interestingly, the yeast p32 homologue has recently been shown to participate in LSU assembly (102); due to its negative charge, it binds several LSU proteins and prevents their aggregation. Thus, it appears conceivable that the effect of *YBEY* KO on the mitochondrial LSU is p32-mediated, and indeed the association of p32 with the LSU was significantly decreased in *YBEY* KO cells (Figure 6G). On the other hand, a heterotrimeric *Trypanosoma brucei* p32 homologue (also known as mt-SAF16/19/25) has recently been identified as part of the mitochondrial SSU “assemblosome” (90). In light of the importance of p32 as a disease-associated and cancer-promoting protein (68,73,74), the discovery of its tight association with YBEY and mitoribosome assembly may guide future mechanistic studies of its so-far elusive molecular functions.

Yet another puzzling aspect is the RNase activity of YBEY. Conserved from bacteria to humans (5,12,19), it is obviously not required for rRNA processing in mitochondria (Figure 4C,D, Supplementary Figure 7). Moreover, substitution H128A, which impairs the RNase activity of the human enzyme, does not affect its ability to complement a *YBEY* KO, similar to the equivalent substitution in *E. coli* (8). Likewise, substitution R55A, which severely impairs the RNase activity of bacterial YbeY (5,12), fully complements *YBEY* deletion in humans and *B. subtilis* (3) and restores 16S rRNA processing in *E. coli* (8). These observations suggest that the RNase activity of YBEY is not necessarily related to ribosome biogenesis. In fact, the promiscuous nature of the human and bacterial enzymes (5,12) is particularly suitable for a turnover RNase. Such a function may be especially important in mammalian mitochondria, which rely on the 3’-exoribonuclease PNPase to degrade RNAs (103) but, like many bacteria, may require an indiscriminate endoribonuclease to initiate decay (104). The previously proposed role of YBEY in ribosome quality control and turnover (5,105) may also be relevant in mitochondria. Further studies are warranted to address these possibilities.

## DATA AVAILABILITY

The RNA-Seq datasets generated during this study are deposited at the Gene Expression Omnibus (NCBI) and accessible through the GEO Series accession number GSE134960. Scripts to reproduce the RNA-Seq analysis are deposited at Zenodo (https:doi.org/10.5281/zenodo.3341191). The mass spectrometry proteomics data have been deposited to the ProteomeXchange Consortium via the PRIDE partner repository with the dataset identifiers PXD014959, PXD015310.

## SUPPLEMENTARY DATA

Supplementary Data are available online.

## ACKNOWLEDGEMENT

The authors wish to thank Soufiane Rhermoul and Timothée Vincent for technical assistance, Valérie Demais and Cathy Royer (Plateforme Imagerie In Vitro, CNRS, UPS 3156, University of Strasbourg) for their help with electron microscopy, Hélène Puccio (IGBMC, Strasbourg) for access to Seahorse, Jérôme Mutterer and Mathieu Erhardt (Microscopy and cellular imaging, IBMP, CNRS, Strasbourg) for their help with FLIM-FRET.

