## supplemental data for "YBEY is an essential biogenesis factor for mitochondrial ribosomes"

**for**

**SUPPLEMENTARY TABLES**

**Table S1. *E. coli* strains used in this study.**

| Strain ID | Background | Plasmids | Resistance | Overexpressed protein gene |
|---|---|---|---|---|
| SAB0021 | BL21 Star(DE3) | pSAP0007 | $Amp^R$ | *$His_6$-MRPL18* |
| SAB0103 | Rosetta | pSAP0077 | $Amp^R$ | *YBEY-$His_6$* |
| SAB0116 | Rosetta | pSAP0079 | $Amp^R$ | *$YBEY^{R55A}$-$His_6$* |
| SAB0117 | Rosetta | pSAP0080 | $Amp^R$ | *$YBEY^{R57A}$-$His_6$* |
| SAB0118 | Rosetta | pSAP0081 | $Amp^R$ | *$YBEY^{D62A}$-$His_6$* |
| SAB0119 | Rosetta | pSAP0082 | $Amp^R$ | *$YBEY^{H118A}$-$His_6$* |
| SAB0120 | Rosetta | pSAP0083 | $Amp^R$ | *$YBEY^{H122A}$-$His_6$* |
| SAB0121 | Rosetta | pSAP0084 | $Amp^R$ | *$YBEY^{H128A}$-$His_6$* |
| SAB0122 | Rosetta | pSAP0085 | $Amp^R$ | *$YBEY^{M137A}$-$His_6$* |
| SAB0123 | Rosetta | pSAP0086 | $Amp^R$ | *$YBEY^{E141A}$-$His_6$* |
| SAB0144 | Rosetta | pSAP0097 | $Kan^R$ | *C1QBP* |
| SAB0146 | Rosetta | pSAP0077, pSAP0097 | $Amp^R$, $Kan^R$ | *YBEY-$His_6$, C1QBP* |
| SAB0199 | Rosetta | pSAP0112 | $Kan^R$ | *MRPS11* |

**Table S2. Human cell lines used in this study.**

| Cell line | Background | Genotype |
|---|---|---|
| Flp-In T-REx 293 | Flp-In T-REx 293 | WT |
| SAL001 | Flp-In T-REx 293 | *FRT::Hyg::YBEY-3×FLAG::FRT::lacZ-Zeo* |
| HepG2 | HepG2 | WT |
| HeLa | HeLa | WT |
| 293T-REx | 293T-REx | WT (*$YBEY^+$*) |
| *$YBEY^+$*_clone1 | 293T-REx | *$YBEY^+$* |
| *$YBEY^+$*_clone2 | 293T-REx | *$YBEY^+$* |
| *$YBEY^+$*_clone3 | 293T-REx | *$YBEY^+$* |
| *YBEY* KO1 | 293T-REx | Δ*YBEY* |
| *YBEY* KO2 | 293T-REx | Δ*YBEY* |
| *YBEY* KO3 | 293T-REx | Δ*YBEY* |
| *YBEY* KO2-YBEY_clone1 | 293T-REx | Δ*YBEY(KO2)::YBEY* |
| *YBEY* KO2-YBEY_clone2 | 293T-REx | Δ*YBEY(KO2)::YBEY* |
| *YBEY* KO2-YBEY-R55A_clone1 | 293T-REx | Δ*YBEY(KO2)::$YBEY^{R55A}$* |
| *YBEY* KO2-YBEY-R55A_clone2 | 293T-REx | Δ*YBEY(KO2)::$YBEY^{R55A}$* |
| *YBEY* KO2-YBEY-H128A_clone1 | 293T-REx | Δ*YBEY(KO2)::$YBEY^{H128A}$* |
| *YBEY* KO2-YBEY-H128A_clone2 | 293T-REx | Δ*YBEY(KO2)::$YBEY^{H128A}$* |
| *YBEY* KO3-YBEY_clone1 | 293T-REx | Δ*YBEY(KO3)::YBEY* |

| | | |
|---|---|---|
| *YBEY* KO3-YBEY_clone2 | 293T-REx | Δ*YBEY(KO3)::YBEY* |
| *YBEY* KO3-YBEY-R55A_clone1 | 293T-REx | Δ*YBEY(KO3)::YBEY*$^{R55A}$ |
| *YBEY* KO3-YBEY-R55A_clone2 | 293T-REx | Δ*YBEY(KO3)::YBEY*$^{R55A}$ |
| *YBEY* KO3-YBEY-H128A_clone1 | 293T-REx | Δ*YBEY(KO3)::YBEY*$^{H128A}$ |
| *YBEY* KO3-YBEY-H128A_clone2 | 293T-REx | Δ*YBEY(KO3)::YBEY*$^{H128A}$ |

**Table S3. Oligonucleotides used in this study.**

**Table S4. Antibodies used in this study.**

| **Antibody** | **Manufacturer** | **Reference** |
|---|---|---|
| Goat polyclonal anti-Actin (C-11) | Santa Cruz Biotechnology | sc-1615 |
| Goat polyclonal anti-ALDH2 (N-14) | Santa Cruz Biotechnology | sc-48838 |
| Mouse monoclonal anti-ATP5A (clone 7H10) | Molecular Probes | A-21350 |
| Mouse monoclonal anti-CYC1(clone 16D10) | Molecular Probes | A-21362 |
| Mouse monoclonal anti-COX1 (clone 1D6E1A8) | Abcam | ab14705 |
| Mouse monoclonal anti-COX2 (clone 12C4) | Molecular Probes | A-6404 |
| Mouse monoclonal anti-COX4 (clone 20E8) | Santa Cruz Biotechnology | sc-58348 |
| Mouse monoclonal anti-cytochrome c (clone 6H2.B4 (RUO)) | BD Biosciences | 556432 |
| Mouse monoclonal anti-FLAG (clone M2) | Sigma-Aldrich | F1804 |
| Mouse monoclonal anti-GAPDH (clone 6C5) | Millipore | MAB374 |
| Mouse monoclonal anti-HDAC2 (clone 3F3) | Santa Cruz Biotechnology | sc-81599 |
| Mouse monoclonal anti-p32 (clone H-9) | Santa Cruz Biotechnology | sc-271200 |
| Mouse monoclonal anti-SDHA (clone 2E3) | Molecular Probes | A-11142 |
| Mouse monoclonal anti-VDAC1 (clone B-6) | Santa Cruz Biotechnology | sc-390996 |
| Rabbit polyclonal anti-bL20m | Sigma-Aldrich | HPA047074 |
| Rabbit polyclonal anti-ERAL1 | Sigma-Aldrich | HPA021425 |
| Rabbit polyclonal anti-HA | Sigma-Aldrich | H6908 |
| Rabbit polyclonal anti-mL38 | This manuscript | 3528 |
| Rabbit polyclonal anti-mS27 | Sigma-Aldrich | HPA071751 |
| Rabbit polyclonal anti-mS35 | Proteintech Group | 16457-1-AP |
| Rabbit polyclonal anti-mS37 | Proteintech Group | 11728-1-AP |
| Rabbit polyclonal anti-mtIF3 | Proteintech Group | 14219-1-AP |
| Rabbit polyclonal anti-NDUFB8 | Sigma-Aldrich | HPA003886 |
| Rabbit polyclonal anti-OPA1 | Proteintech Group | 27733-1-AP |
| Rabbit polyclonal anti-PNPase | Abcam | ab96176 |
| Rabbit polyclonal anti-TOMM20 (FL-145) | Santa Cruz Biotechnology | sc-11415 |

| | | |
|---|---|---|
| Rabbit polyclonal anti-uL22m | Proteintech Group | 16299-1-AP |
| Rabbit polyclonal anti-uL24m | Proteintech Group | 16224-1-AP |
| Rabbit polyclonal anti-uL4m | Proteintech Group | 27484-1-AP |
| Rabbit polyclonal anti-uS11m | Proteintech Group | 17041-1-AP |
| Rabbit polyclonal anti-YBEY | Sigma-Aldrich | HPA018162 |
| Donkey polyclonal anti-Rabbit IgG | Sigma-Aldrich | NA934V |
| Goat polyclonal anti-Rabbit IgG | Agilent | P0448 |
| Goat polyclonal anti-Rabbit IgG | Cell Signaling Technology | 7074 |
| Mouse m-IgGκ BP-HRP | Santa Cruz Biotechnology | sc-516102 |
| Rabbit polyclonal anti-Goat IgG | Sigma-Aldrich | A8919 |
| Rabbit polyclonal anti-Mouse IgG | Agilent | P0260 |
| Sheep polyclonal anti-Mouse IgG | GE Healthcare | NXA931 |
| Goat polyclonal anti-Mouse IgG-Alexa Fluor 488 | Thermo Fisher Scientific | A-11001 |
| Goat polyclonal anti-Mouse IgG-Alexa Fluor 555 | Thermo Fisher Scientific | A32727 |
| Goat polyclonal anti-Mouse IgG-Alexa Fluor 647 | Thermo Fisher Scientific | A-21237 |
| Goat polyclonal anti-Rabbit IgG-Alexa Fluor 488 | Thermo Fisher Scientific | A-11008 |
| Goat polyclonal anti-Rabbit IgG-Alexa Fluor 555 | Thermo Fisher Scientific | A-21429 |
| Goat polyclonal anti-Rabbit IgG-Alexa Fluor 647 | Thermo Fisher Scientific | A-21246 |

**Table S5. Differential gene expression analysis of the mitochondrial transcriptomes of *YBEY*$^{+}$ and *YBEY* KO cells.** Mitochondrial RNAs from three independent cell lines for each genotype were sequenced and the data analysed by DESeq2.

**Table S6. Proteins identified by LC-MS/MS that reproducibly copurified with YBEY-3×FLAG from mitochondrial lysates of HEK293T-REx cells, as compared to the untagged parental control cell line.** Mitocarta2.0 proteins are shown in bold.

**Table S7. Differential quantitative MS analysis of mitoribosomal and mitoribosome-associated/copurifying proteins from three independent *YBEY*$^{+}$ and *YBEY* KO cell lines.** "INF" in the "Fold change" column means an infinite enrichment due to the absence of the corresponding peptides in any of the *YBEY*$^{+}$ replicates. Mitoribosomal and mitoribosome-associated proteins are bold, those significantly underrepresented in *YBEY* KO ($P_{adj}$ < 0.05) are highlighted with yellow, and those depleted >10-fold are red.

**SUPPLEMENTARY FIGURE LEGENDS**

**Figure S1. Occurrence of *YBEY* genes in bacterial and eukaryal clades.** Phylogenetic groups with at least one genome sequenced with contigs of ≥100 kb are shown with the help of the Tree of Life. The presence of identified *YBEY* genes is designated with green dots.

**Figure S2. Mitochondrial localisation of YBEY.** Immunostaining of transiently transfected HeLa and stably transfected HEK293T-REx cells expressing YBEY-3×FLAG. TOMM20 is used as mitochondrial marker. Scale bar, 10 µm.

**Figure S3. Construction of CRISPR-mediated *YBEY* KO cell lines. (A)** Genomic organisation of the *YBEY* locus. The annealing positions of gRNAs and the resulting deletions (Δ) are shown. **(B)** PCR validation of the introduced deletions. WT is the parental HEK293T-REx cell line. Three independent *YBEY*$^{+}$ cell lines were derived from WT as a result of the first (unsuccessful) round of CRISPR/Cas9 mutagenesis. Three *YBEY* KO cell lines were obtained from the corresponding *YBEY*$^{+}$ cell lines after a second transfection that resulted in disruption of all *YBEY* alleles. **(C)** The corresponding western blot confirms the lack of YBEY in *YBEY* KO cell lines.

**Figure S4. *YBEY* KO results in selective deregulation of mitochondrial genes. (A)** Relative mtDNA levels in *YBEY*$^{+}$ and *YBEY* KO cell lines, as assessed by qPCR. Means ± SEM for $n$ = 9 independent cultures from three different cell lines for each genotype are shown; *P*-value, two-tailed Welch's test. **(B)** Snapshot of the mitochondrial *MT-CO1-MT-CO2-MT-ATP8/6* locus showing the downregulation of *MT-CO2* (and to a smaller extent *MT-CO1*) mRNAs in the absence of YBEY. "*8*" designates the small *MT-ATP8* ORF. tRNA genes, poorly covered by the employed RNA-Seq protocol, are not labelled for clarity. **(C,D)** smFISH analysis of *MT-CO1*, *MT-CYB* mRNAs, mt-tRNA$^{Val}$ and 12S rRNA abundance in the *YBEY*$^{+}$ and *YBEY* KO cell lines. Mean total normalised RNA signal area ± SEM is shown in (D) for $n$ = 20 (*YBEY*$^{+}$) and 10 (*YBEY* KO) frames (*MT-CO1*), $n$ = 7 frames (*MT-CYB*), or $n$ = 9 (*YBEY*$^{+}$) and 8 (*YBEY* KO) frames (mt-tRNA$^{Val}$), or $n$ = 8 frames (12S rRNA), respectively; *P*-values, two-tailed Mann-Whitney test. **(E)** Probing total RNA for selected mitochondrial tRNAs by northern blotting reveals no significant differences between *YBEY*$^{+}$ and *YBEY* KO cells. 5S rRNA is used as loading control.

**Figure S5. YBEY is not required for processing or polyadenylation of the various mitochondrial RNA species. (A)** cRT-PCR analysis of the poly(A) tail length of 12S rRNA in *YBEY*$^{+}$ and *YBEY* KO cells. n = 77 plasmid clones from two independent *YBEY*$^{+}$ cell lines and $n$ = 86 clones from two independent *YBEY* KO cell lines. **(B)** RT-qPCR analysis of 4 mt-tRNA precursors with primers spanning 5′ or 3′ tRNA junctions. Means ± SEM for $n$ = 3 independent *YBEY*$^{+}$ and *YBEY* KO cell lines are shown; *P*-values, two-tailed t-test. **(C)** cRT-PCR analysis of the 5′- and 3′-termini of the *MT-CO2* mRNA in *YBEY*$^{+}$ and *YBEY* KO cells. $n$ = 45 (*YBEY*$^{+}$ cells) and 41 (*YBEY* KO) clones are aligned with respect to the reference positions. **(D)** *YBEY*$^{+}$ and *YBEY* KO cells have similar proportions of correctly processed *MT-CO2* mRNA molecules. Proportions and 95% CIs are shown; based on the data in (C); *P*-value, Fisher's exact test. **(E)** cRT-PCR analysis of the poly(A) tail length of the *MT-CO2* mRNA in *YBEY*$^{+}$ and *YBEY* KO cells. $n$ = 45 (*YBEY*$^{+}$ cells) and 41 (*YBEY* KO) clones. **(F)** cRT-PCR analysis of the positions of the 5′- and 3′-termini of 16S rRNA in *YBEY*$^{+}$ and *YBEY* KO cells. $n$ = 100 clones from two independent *YBEY*$^{+}$ cell lines and $n$ = 145 clones from three independent

*YBEY* KO cell lines are aligned with respect to the reference positions. Irregular truncation on 5'-ends of 16S rRNA are particularly frequent in *YBEY* KO cells. **(G)** Both *YBEY*$^+$ and *YBEY* KO cells have high proportions of correctly processed 16S rRNA molecules. However, 16S rRNA is frequently truncated in *YBEY* KO cells. Proportions and 95% CIs are shown; based on the data in (F); *P*-values, Fisher's exact test. **(H)** cRT-PCR analysis of the poly(A) tail length of 16S rRNA in *YBEY*$^+$ and *YBEY* KO cells. $n$ = 100 clones from two independent *YBEY*$^+$ cell lines and $n$ = 145 clones from three independent *YBEY* KO cell lines.

**Figure S6. Human YBEY is a promiscuous endoribonuclease. (A)** Predicted secondary structure of an RNA substrate composed of the 3' minor domain of mitochondrial 12S rRNA (black) and the adjacent tRNA$^{Val}$ (red). The substrate (50 ng) was incubated with increasing concentrations (200, 600, 1000, 2000 nM) of YBEY or the non-nucleolytic control protein uL18m purified under similar conditions, and the cleavage products were analysed by northern blotting with oligonucleotides annealing at the 5'- or the 3'-end of the substrate. The complementary cleavage pattern indicates that YBEY acts as an endoribonuclease. The approximate position of the first major cleavage is shown on the structure with the blue arrow. The orange arrow shows the migration position of mature mt-tRNA$^{Val}$. **(B)** The time-course of the YBEY-mediated cleavage of the precursor of the human mitochondrial tRNA$^{Gln}$.

**Figure S7. Catalytically impaired human YBEY successfully complements *YBEY* KO. (A)** Mutations of conserved residues differentially affect the RNase activity of human YBEY (see also Figure 1A). The 12S rRNA precursor substrate used in (A) was co-incubated with 500 nM purified YBEY proteins and the cleavage products were analysed by northern blotting. **(B)** WT and mutant YBEY versions successfully restore 12S rRNA levels in *YBEY* KO cells. Shown are means ± SEM for normalised 12S rRNA levels measured by RT-qPCR from $n$ = 3 (WT) or $n$ = 6 (*YBEY* KO cell lines) independent replicates; *P*-values, two-tailed Welch's test (all comparisons with the uncomplemented *YBEY* KO cells). **(C)** Metabolic [$^{35}$S]-methionine labelling of mitochondrial translation products reveals that the translational shutdown phenotype of *YBEY* KO cells can be complemented by WT and mutant YBEY alike, i.e., independently of their RNase activity. Western blots for YBEY proteins and VDAC1 and actin as loading controls are shown. All lanes are cropped from the same membrane. **(D)** WT and mutant YBEY restore normal cellular respiration, as assessed by oxygen consumption rate and extracellular acidification measurements by the Seahorse Mito Stress test. Means ± SEM for $n$ = 7 (*YBEY*$^+$ and *YBEY* KO + R55A cell lines), or $n$ = 8 (uncomplemented *YBEY* KO and *YBEY* KO + H128A), or $n$ = 6 (*YBEY* KO + WT YBEY) are shown; *P*-values, two-tailed Welch's test.

**Figure S8. YBEY forms a stable stoichiometric complex with p32 *in vivo*. (A)** Cross-coimmunoprecipitation assays from HEK293T-REx cells that express in either combination YBEY-3×FLAG, p32-HA or untagged proteins with either FLAG- or HA-directed antibodies reveals specific reciprocal copurification of YBEY-3×FLAG and p32-HA proteins. Western blots probed for YBEY-3×FLAG, p32-HA and two non-interacting mitochondrial proteins, mL38 and PNPase, are shown. **(B)** Immunoprecipitation of endogenous p32 from the HEK293T-REx mitochondrial lysate specifically enriches endogenous YBEY, as compared to a control immunoprecipitation from the same lysate with an anti-FLAG antibody. Western blot analysis of lysate (L), flow-through (FT), wash (W) and immunoprecipitate (IP) fractions is shown. **(C)** Analysis of the YBEY:p32 stoichiometry based on $n$ = 6 and 4 coimmunoprecipitation experiments using YBEY and p32 (both endogenous and tagged) as a bait, respectively. Levels of copurified YBEY and p32 were measured by LC-MS/MS and spectral abundance factors for each protein were used to infer their stoichiometry. Medians, interquartile ranges and data ranges are shown.

**Figure S9. YBEY forms a stable stoichiometric complex with p32 in a heterologous system (*E. coli*). (A)** Purification of YBEY-$His_6$ and tagless p32, or both proteins together from native *E. coli* lysates on Ni-agarose columns. Coomassie-stained gels of total lysates (TL), soluble (Sol) and insoluble (Ins) fractions, flow-throughs (FT), washes (W), stepwise elutions with increasing concentrations of imidazole and the beads after the last elution are shown. Only the co-expression of YBEY-$His_6$ with p32 results in a partial solubilisation of the former and specific (at >100 nM imidazole) elution of both proteins. The molecular weight markers are the same as in Figure 5E. **(B)** Analysis of YBEY sedimentation in a high-resolution 10-40% glycerol gradient of the total cell lysate of HEK293T-REx cells. Western blots of YBEY, a small monomeric protein (mtIF3), mitochondrial SSU (mS35) and LSU (uL4m, uL24m, mL38) proteins are shown for comparison. **(C)** Control purification of tagless uS11m from the native *E. coli* lysate confirms that the protein is not retained on the Ni-agarose beads. Designations are the same as in (A).

**Figure S10. *YBEY* KO does not change significantly the *MRPS11* mRNA level.** Means ± SEM for $n$ = 3 independent cell lines are shown; $P$-value, two-tailed t-test.

**Figure S11. Isoelectric point distributions of YBEY proteins from various clades.** Shown are proteobacteria (ancestors of mitochondria), cyanobacteria (ancestors of plastids), major photosynthetic (green) and non-photosynthetic (red) eukaryal phyla. Medians, interquartile ranges, 5$^{th}$ and 95$^{th}$ percentiles and kernel density distributions are provided.

Supplementary Figure 1

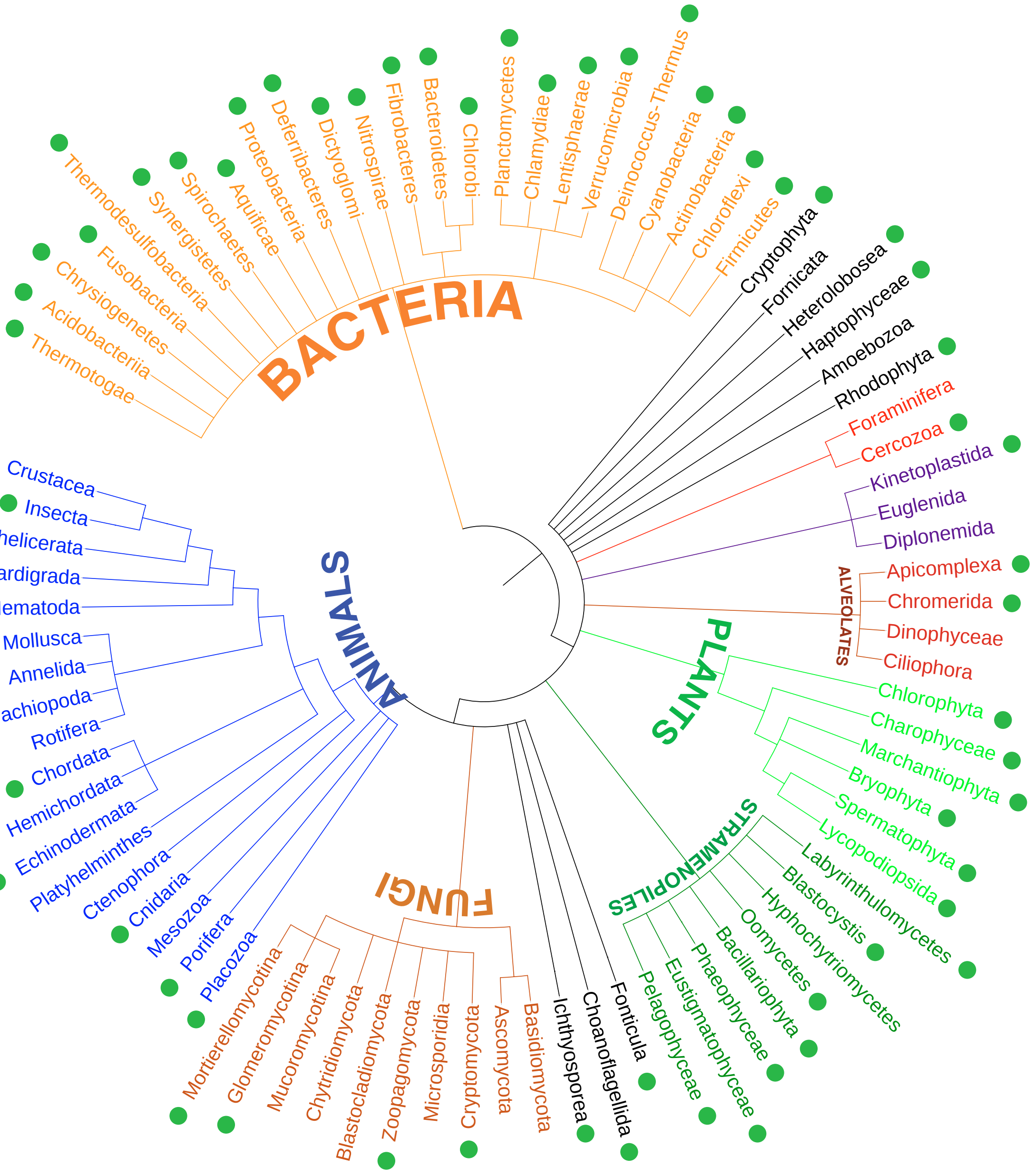

Supplementary Figure 2

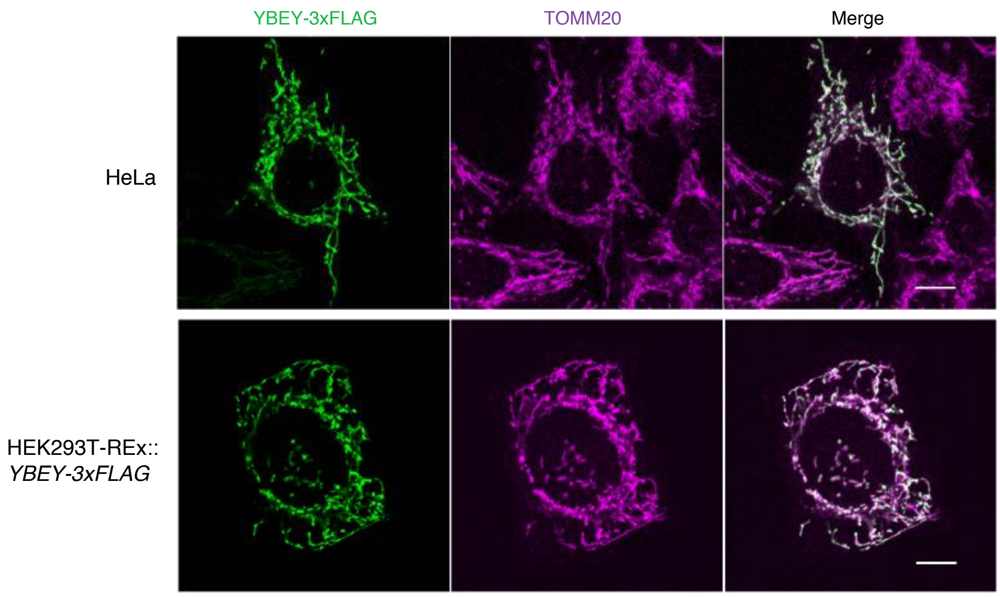

Supplementary Figure 3

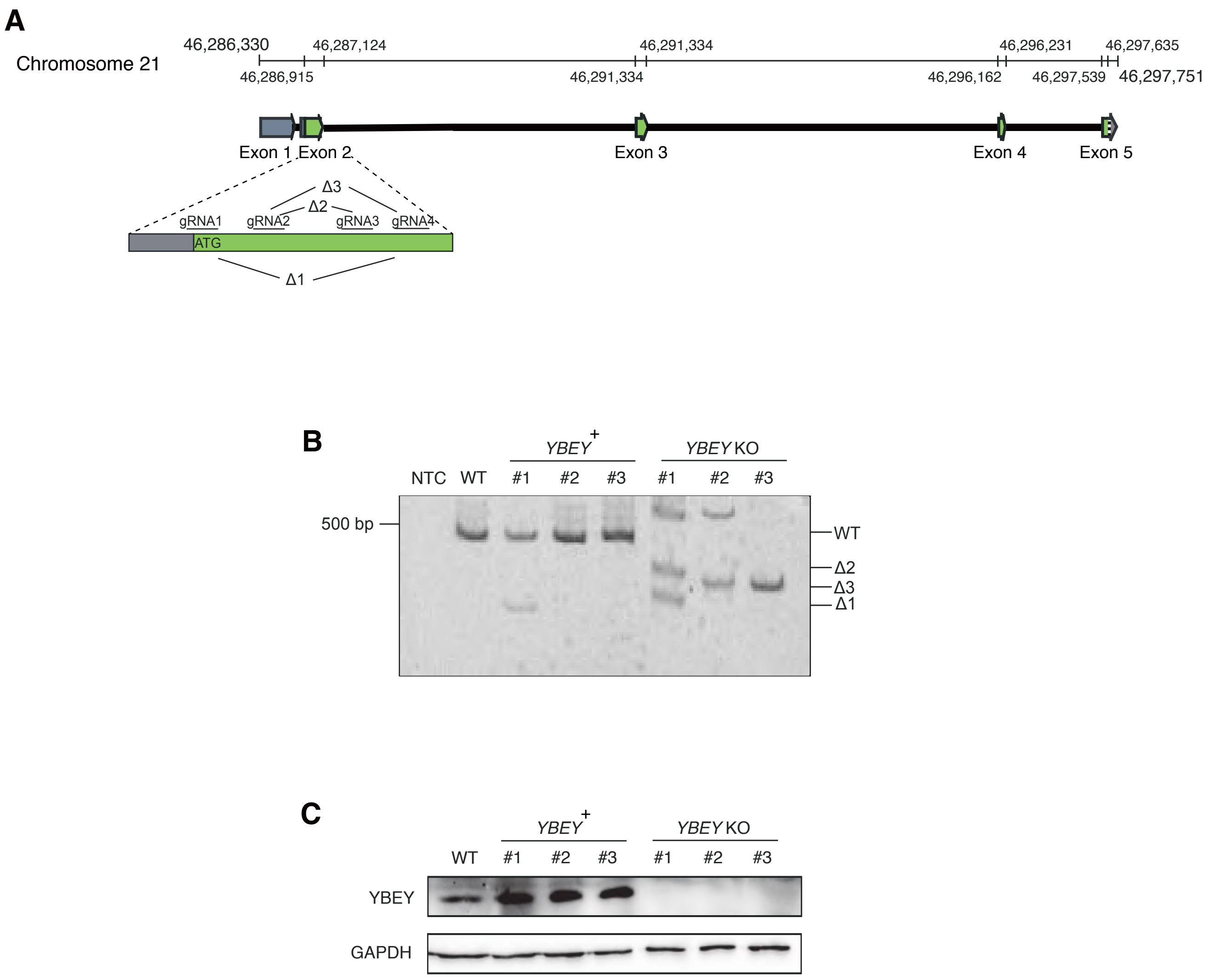

Supplementary Figure 4

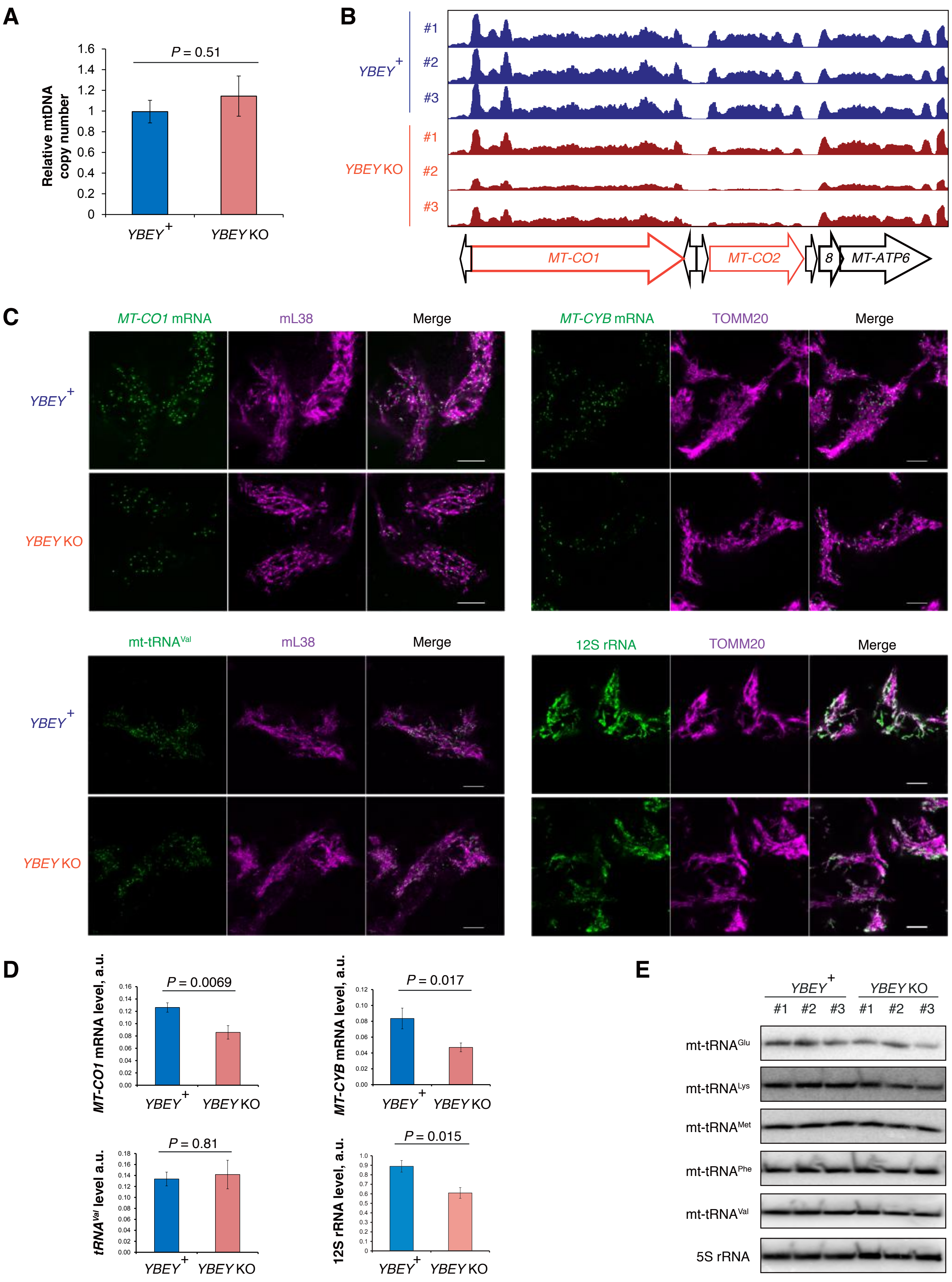

Supplementary Figure 5

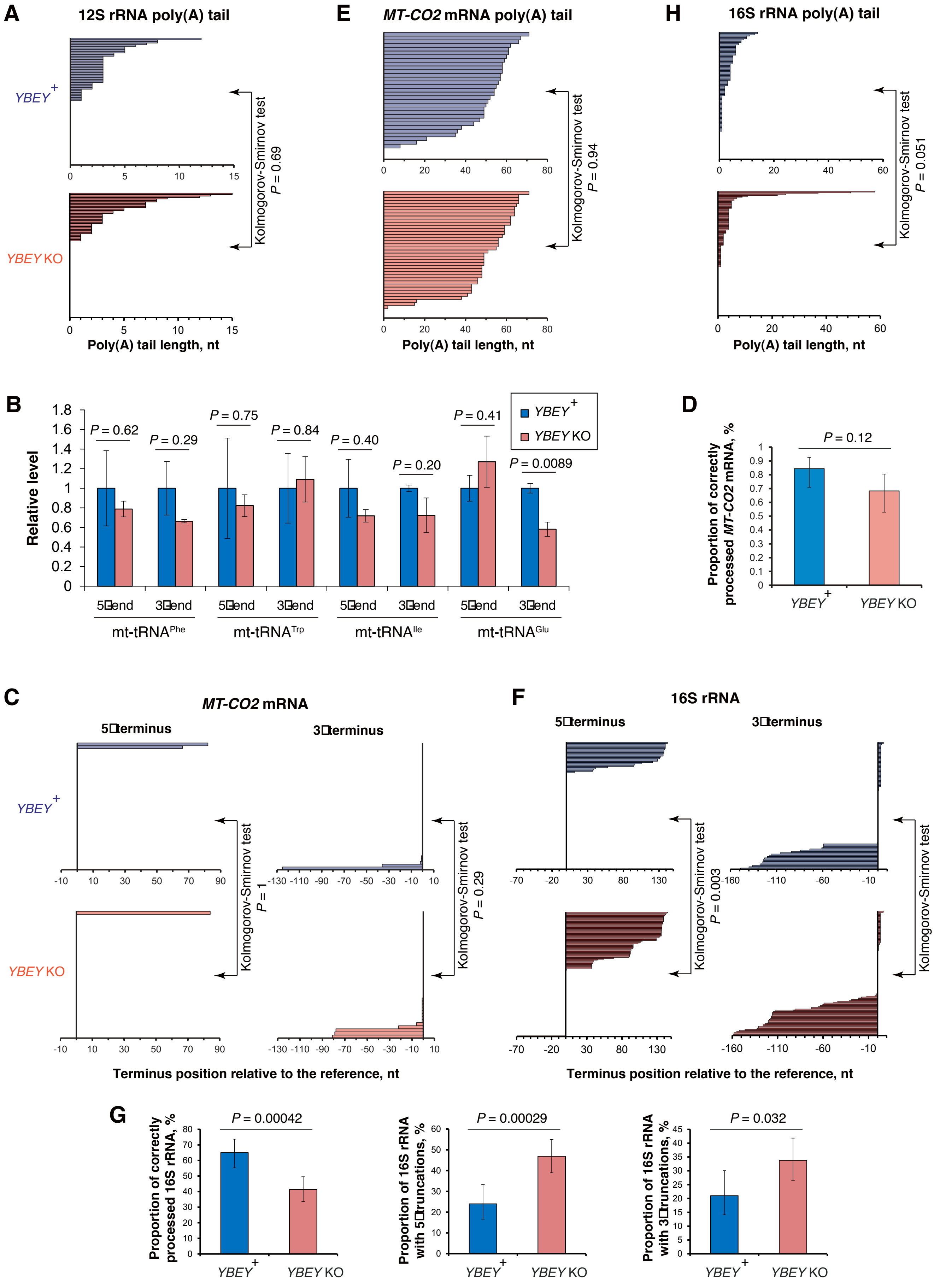

Supplementary Figure 6

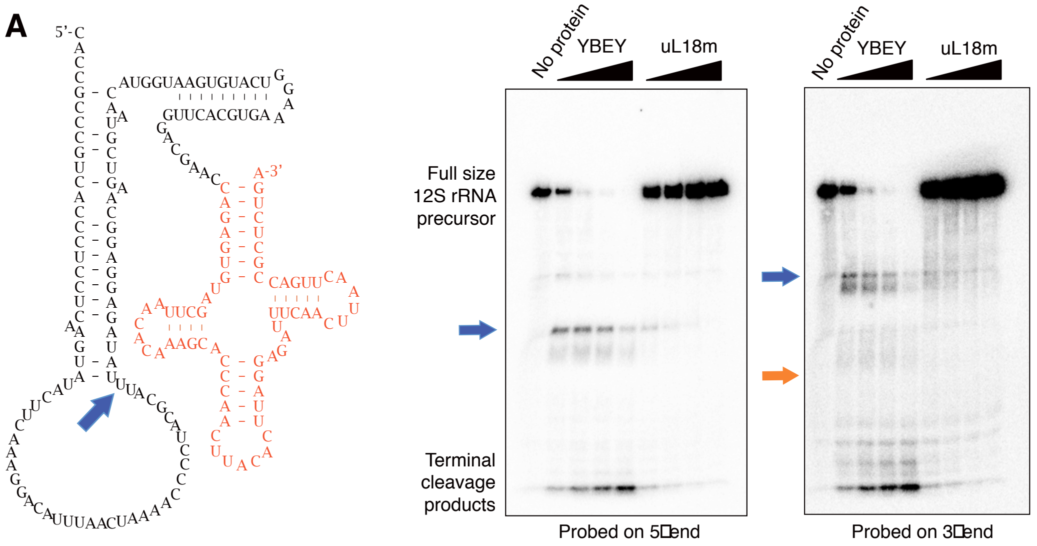


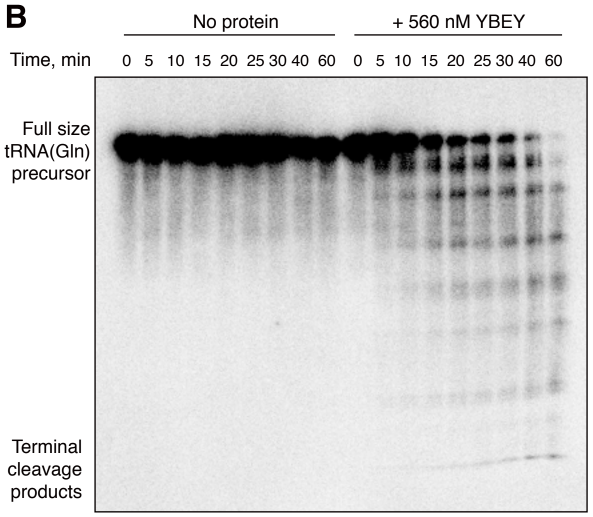

Supplementary Figure 7

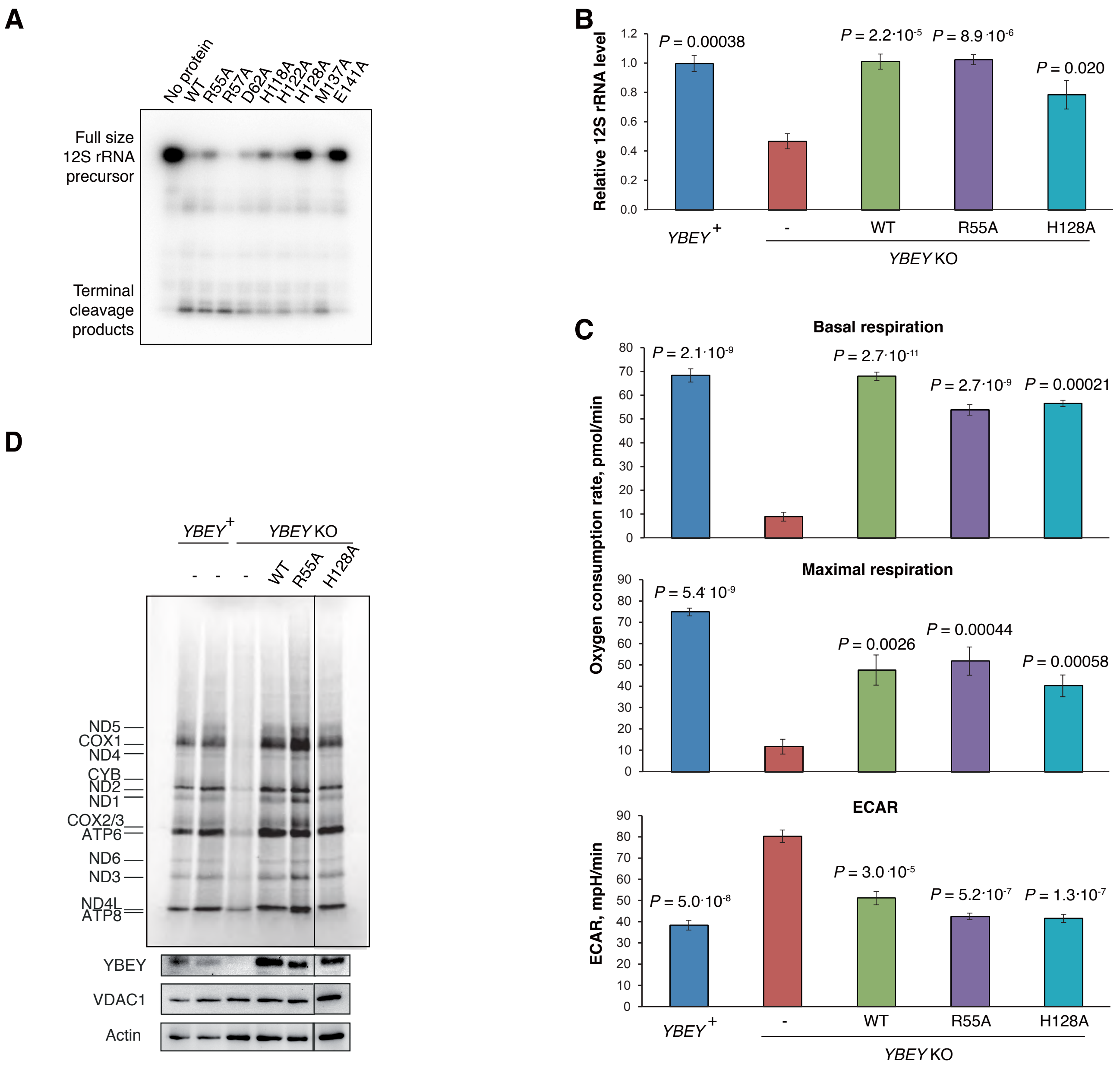

Supplementary Figure 8

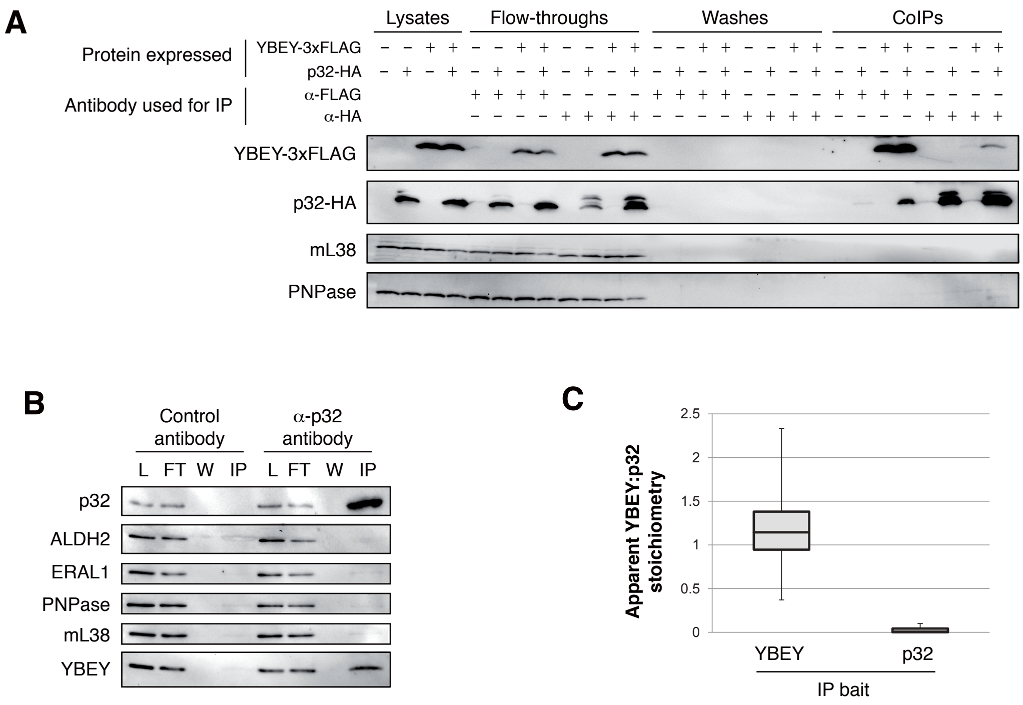

Supplementary Figure 9

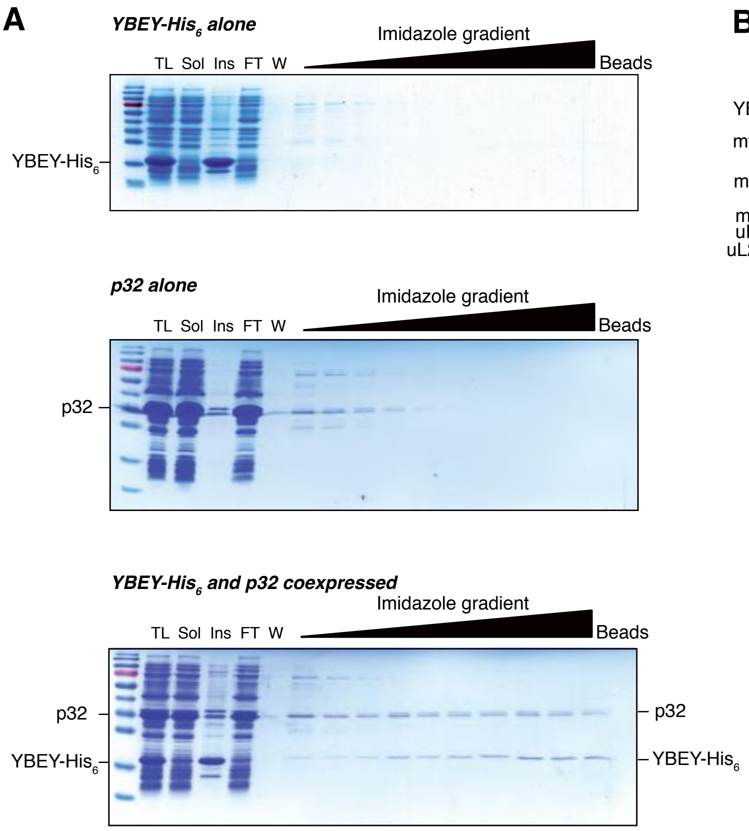


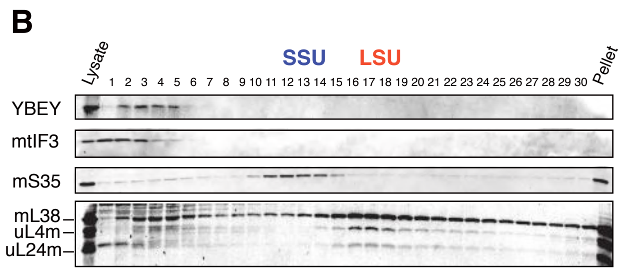


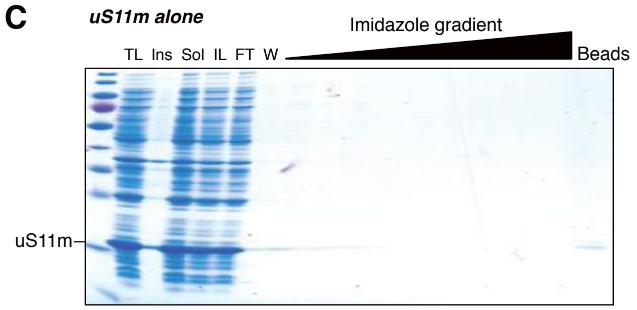

Supplementary Figure 10

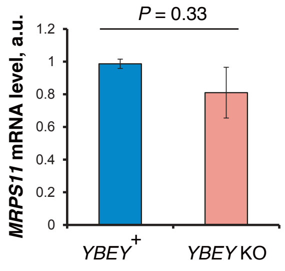

Supplementary Figure 11

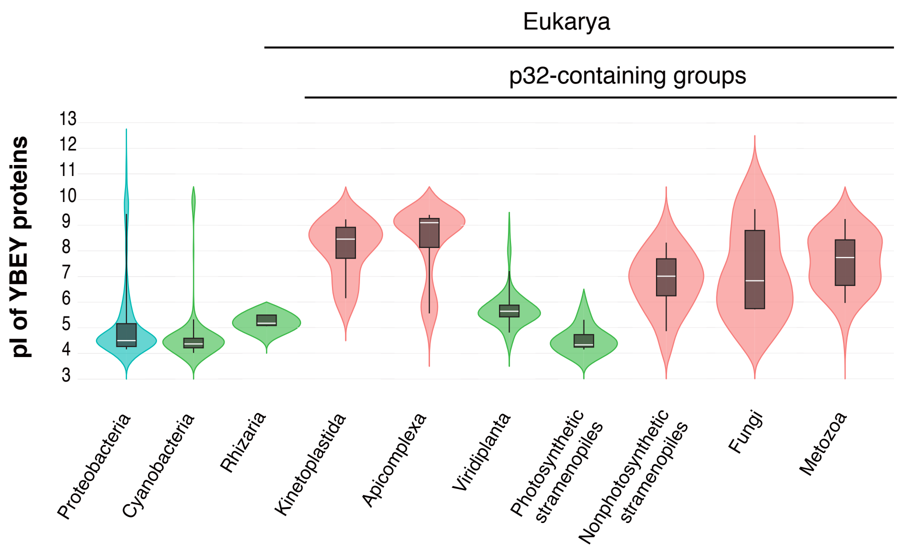
